# Thermal Effects on Neurons During Stimulation of the Brain

**DOI:** 10.1101/2022.04.02.486840

**Authors:** Taeken Kim, Herve Kadji, Andrew J. Whalen, Arian Ashourvan, Eugene Freeman, Shelley I. Fried, Srinivas Tadigadapa, Steven J. Schiff

## Abstract

All artificial stimulation of the brain deposits thermal energy in the brain. This occurs through either Joule heating of the conductors carrying current through electrodes and magnetic coils, or through dissipation of energy in the conductive brain. Similarly, temperature affects all biological processes and chemical reactions. Although electrical interaction with brain tissue is inseparable from thermal effects when electrodes are used, magnetic induction enables us to separate Joule heating from induction effects by contrasting AC and DC driving of magnetic coils using the same energy deposition within the conductors. Since mammalian cortical neurons have no known sensitivity to static magnetic fields, and if there is no evidence of effect on spike timing to oscillating magnetic fields, we can presume that the induced electrical currents within the brain are below the molecular shot noise where any interaction with tissue is purely thermal. In this study, we examined a range of frequencies produced from micromagnetic coils operating below the molecular shot noise threshold for electrical interaction with single neurons. We found that small temperature increases and decreases of 1°C caused consistent transient suppression and excitation of neurons during temperature change. Numerical modeling of the biophysics demonstrated that the Na-K pump, and to a lesser extent the Nernst potential, could account for these transient effects. Such effects are dependent upon compartmental ion fluxes, and the rate of temperature change. A new bifurcation is described in the model dynamics that accounts for the transient suppression and excitation; in addition, we note the remarkable similarity of this bifurcation’s rate dependency with other thermal rate-dependent tipping points in planetary warming dynamics. Furthermore, bifurcations in the steady state dynamics leading to stable firing suppression are described for slightly higher temperatures. These experimental and theoretical findings demonstrate that stimulation of the brain must take into account small thermal effects that are ubiquitously present in electrical and magnetic stimulation. More sophisticated models of electrical current interaction with neurons combined with thermal effects will be required in order to more accurately enable model-based control of neuronal circuitry.

## 1 Introduction

In 1840, J. P. Joule discovered that the “calorific effects of equal quantities of transmitted electricity are proportional to the resistances opposed to its passage”(1). Joule heating is a function of the friction of charge flow within a conductor, and the resistive generation of heat is independent of energy storage within capacitive or inductive circuit elements. Consequently, all stimulation of the nervous system dissipates heat in the electrical components and neural conductors through which electrical current flows. Typically, the effects of neural stimulation are attributed to the induced electrical currents and potential gradients rather than thermal effects. But thermal effects from stimulation are always present.

Investigation of effects of temperature change on neuronal activity was studied by Hodgkin and Huxley (for earlier references see (2)), who observed that the rate of rise and fall of the action potential was greater at higher temperatures (3). Others have reported changes in spike frequency (4), as well as changes in various membrane properties (Table 4). There are thermal effects observed during optogenetic stimulation (5). Recent work has employed infrared lasers to examine a purely thermal effect on neural elements. Such thermal effects have been reported to both stimulate (6; 7; 8) and suppress (9; 10) neuronal activity.

Because magnetic stimulation requires a changing current within a coil to induce an electrical field via Faraday’s law of electromagnetic induction (11), and because mammalian cortical neurons have not been shown to have sensitivity to static magnetic fields (12; 13; 14), magnetic stimulation offers a unique opportunity to disambiguate the effect of thermal dissipation from electrical effects. Such separation is not feasible with electrical stimulation where the effects of Joule heating and electrical current modulation of neuronal activity are inseparable.

Magnetic neuromodulation techniques utilizing micro-coils have garnered significant interest in recent years. Small size improves spatial selectivity as well as enabling implantation (15). Interestingly, both activation (16; 17; 18; 19) and suppression (20; 21) of neuronal activity have been experimentally demonstrated, and theoretical explanations for magnetic activation of individual neurons have been proposed (22; 23). Nevertheless, tissue near the coils will experience temperature increases from the Joule heating of both the coils and from inductively driven current flow within tissue (24).

We here present a set of single neuron experimental measurements where we controlled for the Joule heating within micromagnetic coils using equivalent power dissipation from AC and DC driving currents, while monitoring the temperature changes near the targeted neuron. Consistent suppression and rebound excitation effects were observed from the thermal effects, and a computational and theoretical framework constructed to explain the biophysics of this phenomenon. These thermal effect findings are applicable to all types of stimulation of the brain.

## 2 Results

### 2.1 Experimental thermal effects on spike height and frequency

We observed the activities of cortical layer V pyramidal cells from male Sprague-Dawley rat brain slices *in vitro*, using a loose-patch attachment (25) to reduce the possibility of electrical shunting into the neuron through the electrode during magnetic stimulation as in (26). Stimulation was applied by driving micromagnetic coils with either AC or DC current adjusted for equivalent power dissipation (*P* = *I*^2^*R*) and thus equivalent temperature change measured near the patched cell (Fig. 1a,b see *Methods*). Unexpectedly, we observed that the pattern of spike rate changes with AC or DC stimulation could be identical, with transient suppression at the onset of stimulation current and transient hyperexcitability after removal of current, accompanied by similar changes in temperature at the level of the coils and cells (Fig. 1c,d).

**Figure 1:**
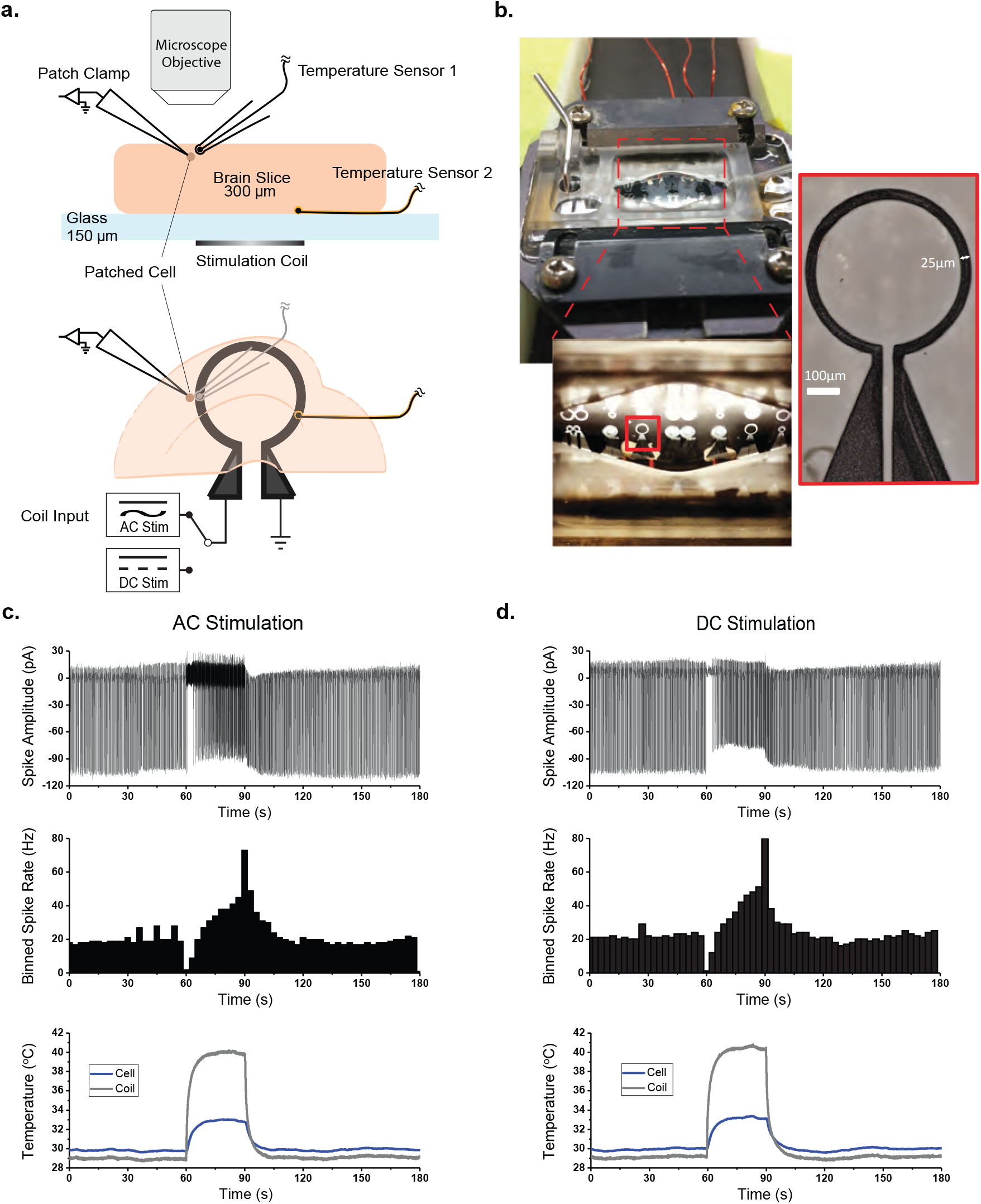
Experimental setup and representative traces. **a**, Depth (above) and overhead (below) view schematic of experimental setup, showing the location of the patch clamp recording electrode, temperature sensors, and the stimulation coil relative to the brain slice. **b**, Photo of the recording chamber and stimulation coils. **c**, Representative responses of layer V pyramidal cells to AC stimulation (500 Hz continuous sine, left column) and DC stimulation (right column), binned spike rate over the stimulation trial (middle) and temperature measured near the patched cell and coil (bottom).

We reduced the amplitude of the stimulation current in the micromagnetic coils to limit the temperature increase near the patched cell to about 1°C for 180 seconds, and then allowed adiabatic cooling back to the 30°C bath temperature baseline upon cessation of coil stimulation (Fig. 2d). We verified this equivalent temperature increase by placing a temperature sensor near the patched cell (Fig. 1a), and found the time constant of temperature increase and decrease to be on the order of 5 seconds (Fig. 2d).

**Figure 2:**
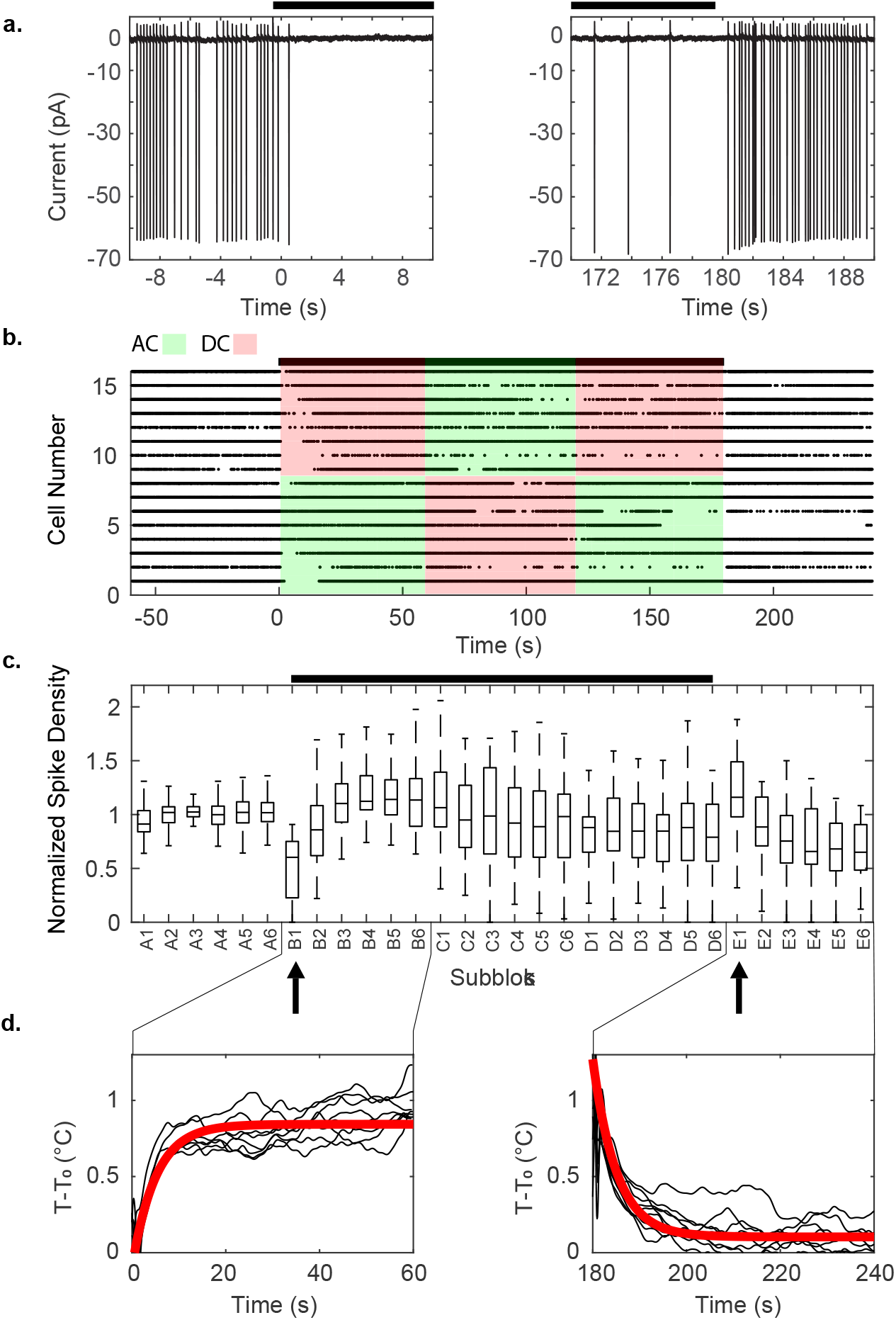
Example of experimental block design limiting heating to 1°C from the highest frequency trials at 200 kHz. This very high frequency would be unlikely to modulate the much slower spike generating mechanisms. The black bars indicate when stimulation was applied (AC or DC). (**a**) Individual spikes observed in the experiment, near the times where the warming and cooling began. Since the cell is patched in voltage-clamp mode, the measurements are in current units, and depolarization of membrane potential results in negative input current. (**b**) Raster plot of spikes from the 200 kHz stimulation experiments. (**c**) Boxplot of normalized spike density from control and stimulated blocks of all experiments. A 180-second stimulation period consisted of three blocks of alternating AC and DC current stimulation, inserted between two 60-second control blocks, for a total of five 60-second blocks labeled A to E. We further subdivided each block into six equal-length sub-blocks of 10 seconds each, as A1-A6, etc., and compared the spike densities and heights across all sub-blocks. Both ANOVA and Kruskal-Wallis tests with subsequent Tukey-Kramer post hoc analysis showed significant suppression of spikes immediately following the application of heat at sub-block B1 (left arrow, ANOVA *df* = 63, *F* = 69, *p* < 10^-19^; Kruskal-Wallis *p* < 10^-7^), and hyperactivity following the removal of heat at sub-block E1 (right arrow, ANOVA *df* = 63, *F* = 3.1, *p* < 0.032; Kruskal-Wallis *p* < 0.009). These changes in spiking activity were transient. We detected no statistically significant changes in the height of the spikes. (**d**) temperature measurements, and the line of best fit. The time constants for heating and cooling were 5.048 and 5.033 seconds with *R*^2^ = 0.6940 and *R*^2^ = 0.5866 respectively.

During AC stimulation the coil was driven with continuous sinusoids to induce changing magnetic fields at 50 Hz, 500 Hz, 5 kHz and 200 kHz, while controlling the current amplitude to maintain the same power dissipation and 1^°^C temperature increase. In order to isolate thermal effects from magnetic stimulation, we delivered AC and DC current delivered in a block design, alternating AC-DC-AC with DC-AC-DC stimulation blocks with flanking control blocks without stimulation (Fig. 2b). Each stimulation block was subdivided into 10 second sub-blocks (per 60 second block). This enabled us to determine whether any magnetically driven electrical current induction affected the cell’s action potential frequency or height within each sub-block.

We identified a statistically significant transient suppression of spikes immediately following the onset of stimulation, and transient hyperactivity following cessation of stimulation (Fig. 2a,c). This effect was independent of whether AC or DC stimulation was applied to the coils, and the equivalent power dissipation was reflected in identical temperature increase and decrease profiles (Fig. 2d, see also supplementary Fig. S1).

Across all AC stimulation frequencies, we found no statistically significant differences in spike height or spike frequency within a given sub-block (Table S2). That is, the changes seen at the onset and offset of stimulation appeared independent of driving frequency applied.

### 2.2 Detecting the effects of AC stimulation on spike timing

We next determined whether spike timing was changed by the AC stimulation sinusoids. This was done by testing whether the spikes tended to occur at particular phases of the AC sinusoids. The statistical distributions of the phases of the spike times were evaluated using the Raleigh Z statistic for circular distributions (see *Methods*). No AC blocks showed statistically significant spike phase preference compared to the DC blocks (Fig. S6). We also measured the directionality of spike phases in 3 second time windows to test whether spike entrainment might occur more often in AC blocks than in the DC blocks over short periods of time. To reduce the chance of type II error, we identified candidate time windows showing entrainment by comparing data with randomly shuffled surrogate interspike intervals drawn from the same data blocks and analyzing the spectral power density of the ensemble (see *Methods*). The populations of Rayleigh Z p-values calculated from these candidate time windows for AC and DC blocks were compared to each other using the Anderson Darling test (27) and Wilcoxon ranksum test. The results (Table 1) showed that there was no statistically significant difference between the spikes in DC and all frequencies of AC blocks. This extensive set of the statistical analysis of experiments and controls was required to prove that the effects on neural activity were due to thermal effects rather than any detectable trace of magnetic induction of electrical currents interacting with the neurons.

**Table 1:**
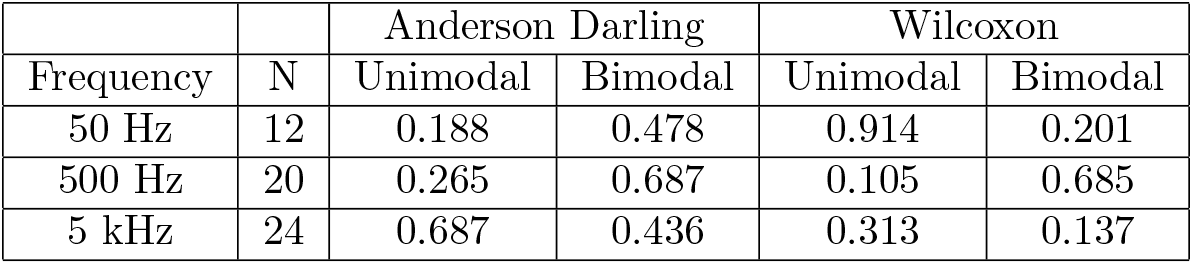
P-values obtained from comparing the distributions of the Rayliegh Z p-values in AC and DC blocks. Since all tests at all paradigms and frequencies failed to reject the null hypothesis, we did not detect any evidence of spike entrainment to the AC stimulation phase. N is the number of measurements taken. Since each cell was measured twice in the two different stimulation paradigms (AC-DC-AC and DC-AC-DC), N represents two times the number of cells recorded. Experiments done at 200 kHz are not applicable to this analysis of spike timing relative to the phase of stimulation.

### 2.3 Modeling temperature-mediated transients

To understand the thermal effects observed in our experiments we modified a computational model based on (28), which extends the original Hodgkin-Huxley formalism (29) to include relevant structural micro-anatomy, conservation of charge and mass, energy balance, and volume changes. The model consists of a single neuronal compartment surrounded by a thin extracellular space that is connected to the bath *in vitro* (or to a capillary *in vivo*) through ionic diffusion (Fig. S2). These biophysical components are required to unify a range of phenomena such as spikes, seizures and spreading depression, which the original Hodgkin-Huxley framework does not characterize well for mammalian neurons. The Nernst potentials that underlie the transmembrane voltage dependency on ionic concentration differences are proportional to temperature. In addition to the thermal effect inherent in the Nernst potentials, we identified eleven membrane processes in the model that could be important in explaining the thermal effects observed: maximal Na-K pump rate, maximal *Na*^+^ and *K*^+^ channel conductances, rate of gate dynamics (*dn/dt, dm/dt, dh/dt*), membrane leak conductances for *Na*^+^, *K*^+^ and *Cl*^−^, and strength of cotransporters NKCC1 and KCC2. The effects of these thermally sensitive processes on membrane potential can be categorized as hyperpolarizing (pump strength, *K*^+^ and *Cl*^−^ conductances, and Nernst potentials), depolarizing (*Na*^+^ conductances), or charge-neutral (gating kinetics and cotransporter strengths).

Although biological processes have a complex relationship to temperature (30), individual cellular processes generally accelerate when temperature is raised a small amount. We used *Q*_10_ factors to model these accelerated rates. The *Q*_10_ factor of a process *y*, describes the change of reaction rate when the temperature is raised by 10°C. For an arbitrary temperature change, *T – T*_0_, the new maximal rate of a process *y*(*T*), with initial process rate *y*(*T*_0_) is calculated as:

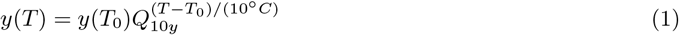

When all temperature dependencies in the model are used to simulate the effect of temperature increase or decrease from baseline, transient spike suppression or hyperactivity is observed as in the experiments (Fig. 3a-d). We next examined the effect of individual model components on cellular activity. Increasing the temperature-mediated effects of the hyperpolarizing processes transiently silences the cell, while increasing the effects of the depolarizing processes induces hyperactivity (Fig. S3). Decreasing the temperature-mediated effects of these parameters has opposite results, while changing charge-neutral parameters does not create significant transient behaviors. Since our *in vitro* data show silencing at the application of heat and hyperactivity at the removal (Fig. 2a), the hyperpolarizing components contained in the model appear to dominate the cell’s thermal response.

**Figure 3:**
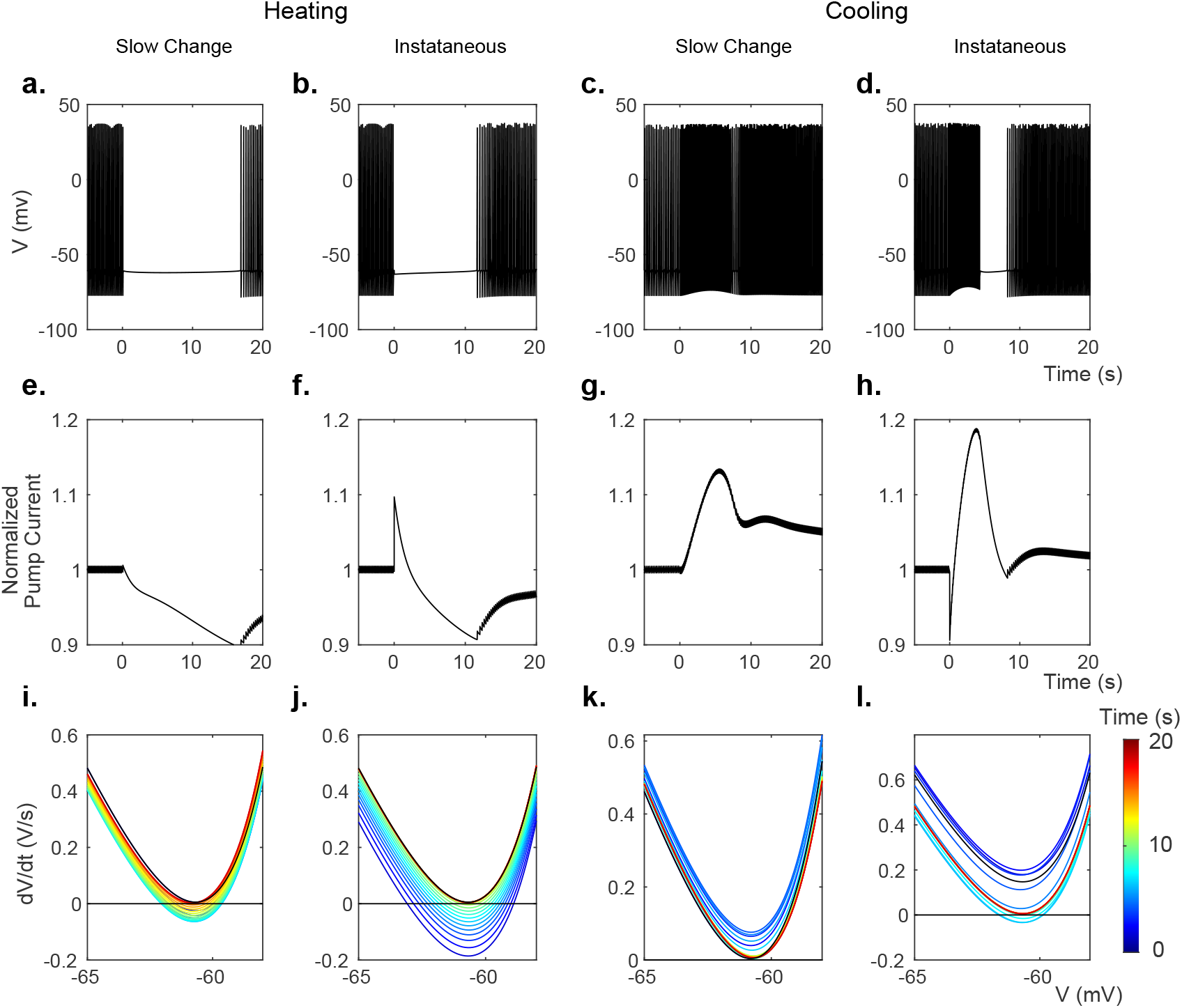
Modeled membrane voltage *V* (a,b,c,d), normalized Na-K pump current (e,f,g,h), and rate of change of voltage 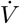 (i,j,k,l) during transient periods. **a,e,i** and **c,g,k** were modeled while changing all membrane parameters slowly with a 5 second time constant to their values at 1° C temperature increase, using *Q*_10_ values from Table 5. **b,f,j** and **d,h,i** were modeled while changing only the maximal pump rate instantaneously to or from 110% of baseline. All times are measured from the onset of temperature change.

We next use the simplifying assumption that similar membrane process categories (ion transport, Nernst potentials, etc.) will have similar *Q*_10_ factors (Table 4). When the *Q*_10_ factors are the same, the cellular process rates will change by the same proportion for any change in temperature. We simulated a temperature rise as a step increase of temperature affecting each grouped category of membrane processes (hyperpolarizing or depolarizing) by 105%, 110%, and 140% followed by a return to baseline values (Fig. S4). Changing either the leak conductances or the voltage-dependent channel conductances induced hyperactivity at the onset of heating, rather than silencing as observed in the experiments. Thus, under assumptions of similar *Q*_10_ factors, the depolarizing *Na*^+^ currents would dominate, and this is inconsistent with the experimental findings. Relaxing the *Q*_10_ similarity constraint for similar biophysical processes, such as leak currents, still reveals that *G_NaLeak_* would dominate the hyperpolarizing effects of *G_KLeak_* and *G_ClLeak_* for similar temperature increases (Fig. S5).

The voltage gated channel conductances, *G_K_* and *G_Na_*, do not substantially suppress the cell activity during heating even when their values are set to maximal (Fig. S3). In contrast, the Na-K pump current demonstrates transient suppression with a temperature increase, and transient hyperactivity with temperature return to baseline (Fig. S3). In addition, we found that increasing the temperature in the Nernst potential equations for Na, K, and Cl gave a similar transient suppression and hyperactivity as the Na-K pump (Fig. S3). The charge-neutral co-transporters had no effect on model cellular activity.

### 2.4 Ion concentrations characterize a temperature sensitive bifurcation

When the model is periodically firing, without thermal perturbation, it is in a stable limit cycle. This model contains variables characterized by fast and slow time scales (Fig. 3a-l). The fast variables are the membrane voltage *V* and the three gating variables *m, n*, and *h*. The slow variables are the analyte concentrations inside and outside the cell. These concentrations define the environment in which action potentials are generated, establish the Nernst potentials and set the pump currents, thereby modifying the spiking behavior. When a dynamical system in a stable limit cycle is perturbed by an abrupt change of parameters, such as a step change in temperature, the system must then traverse through the state space from the old stable state to the new one (provided the new parameters also define a new stable basin of attraction).

There are substantial qualitative differences in spiking activity when temperature is changed slow or fast. The spiking behavior of the model in Fig. 4a,b demonstrates the effect of the speed of the temperature change on the model cell’s spiking. Faster temperature time constants create transient silencing and hyperactivity in the model analogous to the temperature effect on *in vitro* neurons in Fig. 1c,d and 2a. In Fig. 4c,d, the model trajectories resulting from faster rates of temperature changes take large detours in state space, rather than the more direct paths of slower changes. Slow changes in temperature do not alter the equilibrium of the system much, and allow the faster variables to come to equilibrium at all times adiabatically. Slow changes in temperature can even preserve the stable limit cycle, shown as broad ribbons within the trajectories in state space reflecting the oscillations of the spiking limit cycles (time constants of 50 and 500 s in Fig. 4). With slower temperature changes the model does not display transient silencing or hyperactivity. But for fast temperature changes, the new temperature driven equilibrium can be far away from the present state, and needs to be reached as the system readjusts by its own internal fast and slow dynamics. Such substantial trajectory excursions through state space, arriving at the same final equilibrium point, may take the system far from its original limit cycle frequency and may abolish the spiking entirely as shown in the model and experiment.

**Figure 4:**
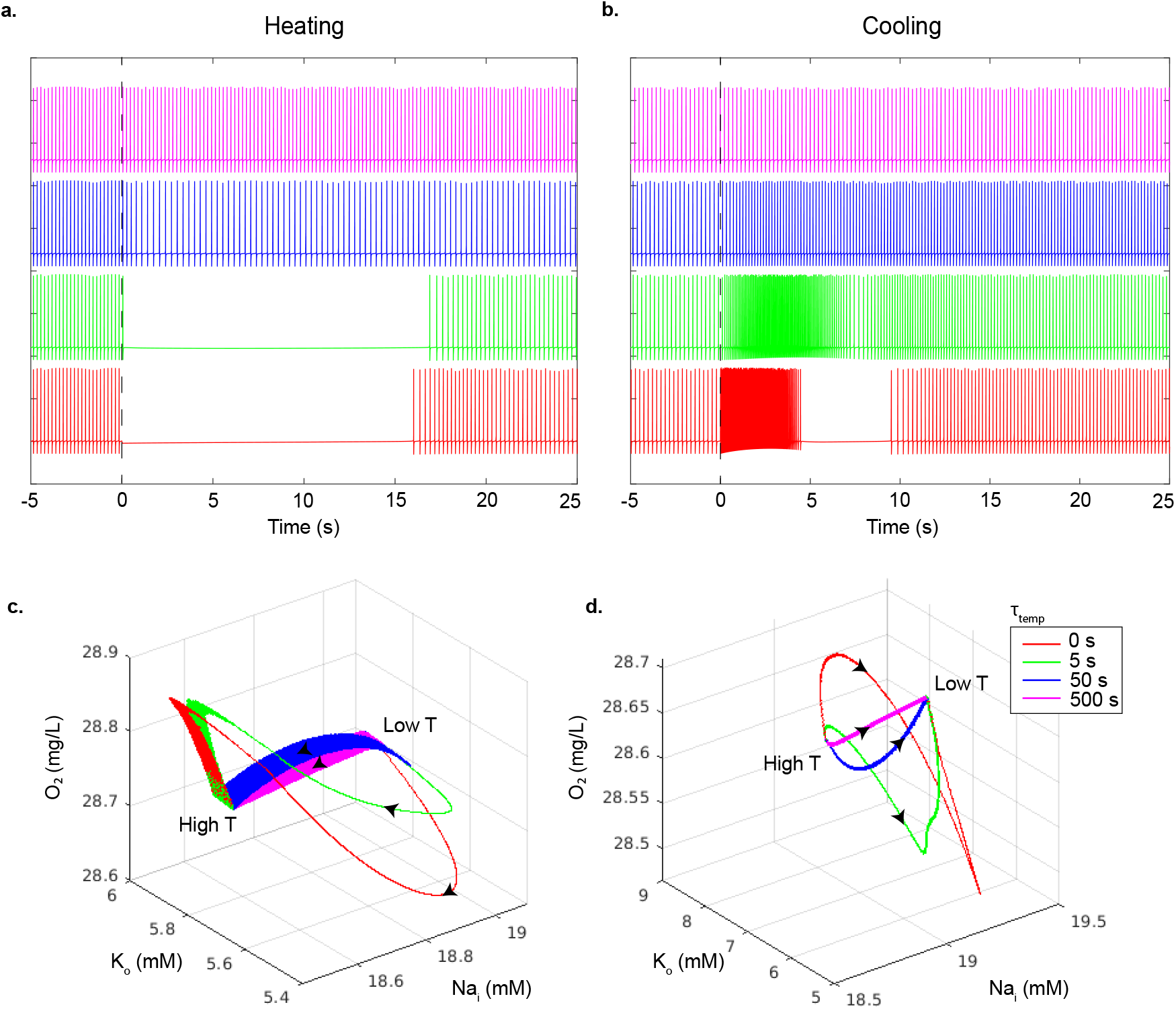
Modeled membrane voltage as a function of the rate of the temperature change. All membrane parameters were changed with decreasing time constants (500, 50, 5 and 0 seconds, top to bottom respectively) to their values for a 1°C temperature increase in **a**, and a 1°C decrease in **b**, using the *Q*_10_ values in Table 5. The onset of temperature changes occur at time zero in both plots. The bottom row shows the model neuron dynamics in the pump state space (*O*_2_, *K_o_, N_a_i__*) as a function of how fast the temperature change is applied for warming (**c**) and cooling (**d**). Black arrows represent the direction that the state space trajectories traverse for warming (temperature low to high, **c**), and cooling (temperature high to low, **d**).

These transient effects can be described by a type of bifurcation whose stability is characterized as a saddle-node. First, we separate the model state vector into fast and slow variables. Then we treat the trajectory of the slow variable state vector ***C***(*t_s_*) = [[*Na*^+^]*_i_*, [*Na*^+^]*_o_*, [*K*^+^]*_i_*, [*K*^+^]*_o_*, [*Cl*^−^]*_o_*, [*O*_2_]] as a function of slow time *t_s_*. We also assume that the gating variables with their fast time scales are at their steady state values. Then, we plot 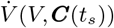 vs *V* (Fig. 3i-l). The x-intercepts of this graph are the equilibria of V. We can understand how temperature affects the spiking behavior of the model by studying these fixed points. At the onset of a modeled temperature increase (Fig. 3i,j), there exists one stable equilibrium point (P1) at −63.0 mV and two unstable equilibria points (P2 and P3) at −58.9 mV and −41.1 mV (see supplementary Fig. S7). The stable fixed point at P1 corresponds to quiescent cell behavior as seen in the *in vitro* patch-clamp data in Fig. 2a,b, where the cell ceases firing upon warming.

We determine the stability of the gating variables at the equilibria by calculating the eigenvalues of the Jacobian of *V* and the gating variables m, h, and n at each equilibrium. The Jacobian and its components are shown in section 6.2. All eigenvalues must have negative real parts for the equilibirum to be stable. This analysis shows that P1 is the only stable equlibirum.

We can verify this result near these equilibrium voltages, where we know that *dm/dt* >> *dn/dt* >> *dh/dt* (Fig. S8). To illustrate the stability of the equilibrium points, we generated field maps for each of the gating variables versus *V*, assuming that for each gating variable, faster gating variables are at their steady state values for the corresponding voltage (*x*_∞_(*V*) = *α_x_*(1 – *x*) – *β_x_*, where *x* = *m, n, h*), and slower gating variables are at their steady state values for the equilibrium voltage (*x*_∞_(*V_eq_*)) in Fig. S9 (31). For example, in calculating the field map of n vs V, we assumed that m will be at *m*_∞_(*V*), and h will be at *h*_∞_(*V_eq_*). This analysis confirmed that P1 is the only stable equilibirum.

Regarding the slow variables, we treat the slow time, *t_s_*, as the bifurcation parameter for the equilibria. As *t_s_* progresses following warming, and by extension the analyte concentrations shift, the 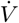 curve moves up. Eventually the two equilibria (points P1 and P2) collide, creating a saddle-node bifurcation. Once the stable equilibrium disappears (the quiescent state), the cell is able to depolarize further to initiate action potentials. In a similiar manner the 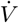 curve can also be used to predict hyperactivity upon cooling. Fig. 3k,l show the 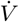 curve when the heat is removed. The distance between the local minimum of the curve and the line 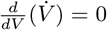 is representative of the cell’s depolarization speed, and consequently, the frequency of spikes. When the curve is below this line, the distance between the curve minimum and 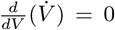 represents how resistant the cell is to spiking.

### 2.5 Further Effects at Equilibrium

To further characterize the biophysics, we reduced the complexity of the ion species relationships (see *Methods*) and used numerical continuation analysis to explore bifurcations at a broader range of temperatures. While models of small changes in temperature showed transient changes but not qualitative differences in longer term equilibrium spiking behavior, bifurcation analysis of the reduced model at higher temperature revealed a subcritical Hopf bifurcation around 37.26^°^C (Fig. 5a). Creation of this bifurcation only requires the Nernst potential to change with temperature; it exists even when all other *Q*_10_’s are set to 1. When the temperature dependence of the Nernst potential is introduced to the basic HH model without any charge or ion conservation, the Hopf bifurcation point changes to a saddle-node, but the stable branch at high temperature still emerged.

**Figure 5:**
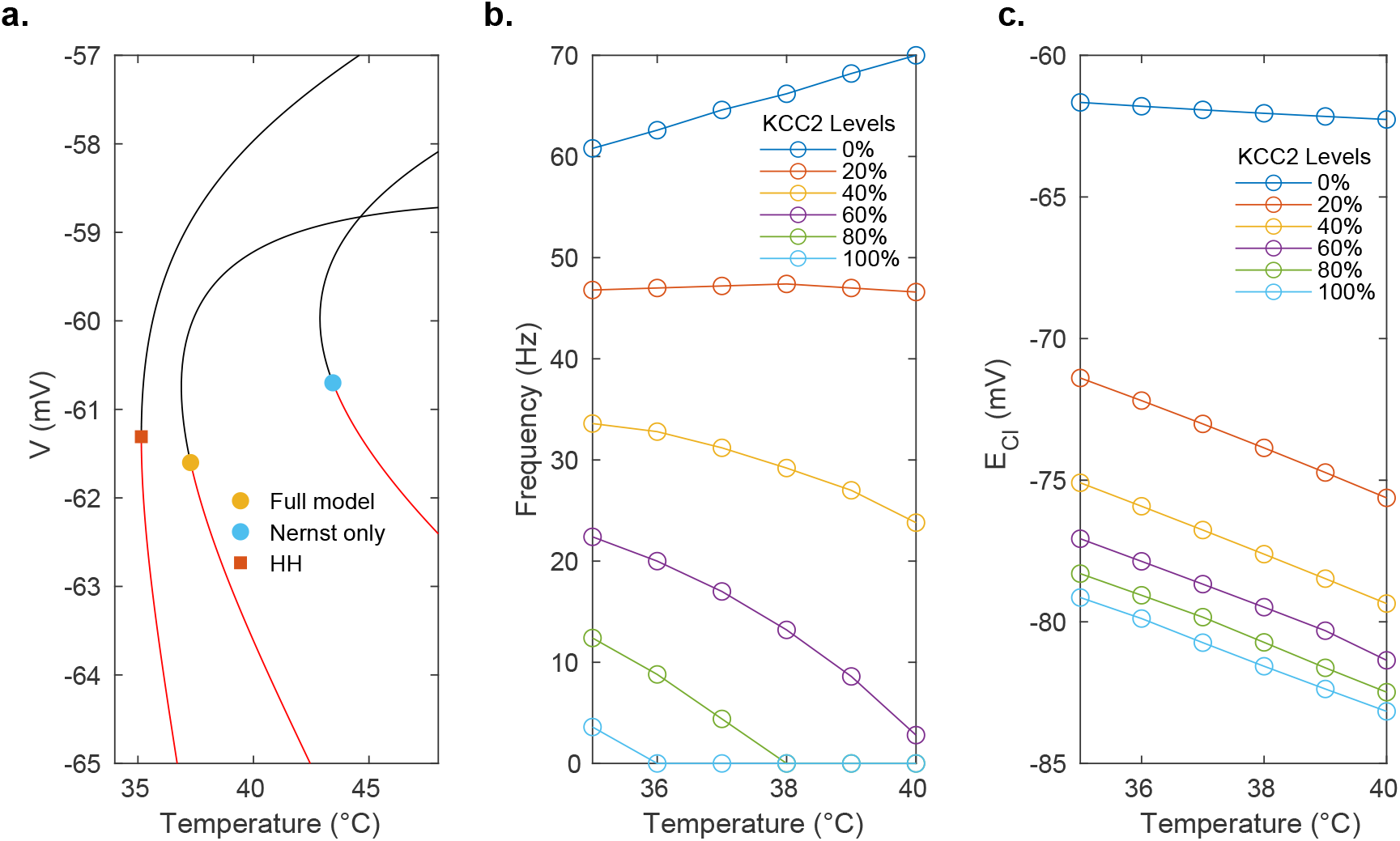
Equilibrium temperature effects. **a**, Bifurcation diagrams for different models. Red lines are the stable equilibria, and black lines are unstable equilibria. The square symbol represents a saddle-node bifurcation, whereas the circle symbols represent subcritical Hopf bifurcation points. The Nernst only model sets all *Q*_10_ values set to 1, and HH is the original Hodgkin-Huxley model with no ion conservation. The nature of the bifurcation points was determined numerically. **b**, Spiking frequency as a function of KCC2 transporter expression and temperature. **c**, Chloride reversal potential, *E_Cl_*, as a function of KCC2 expression levels and temperature.

While all membrane parameters affect the location of this bifurcation point, the KCC2 cotransporter expression level also qualitatively changes the behavior of the cell as the temperature is increased. In the less mature developing brain, it is known that KCC2 levels are not as high as in the mature brain, and febrile seizures are more commonly observed with hyperthermia (typically at the onset of fever as temperature is increasing). In the full model with normal mature KCC2 levels (100% strength in our model), spiking is suppressed above 36°C. At normal KCC2 levels, the frequency and temperature describe an inverse relationship in the full model, while at lower KCC2 levels, the relationship becomes positively correlated and spiking is not suppressed (Fig. 5b). Furthermore, the equilibrium chloride level, *E_Cl_*, becomes elevated (more depolarized) at lower levels of KCC2 expression, reducing the effectiveness of inhibitory transmitter on the cell (Fig. 5c). These effects of KCC2 combine to render spiking at elevated temperatures more prominent in the immature brain with lower levels of KCC2 expression.

## 3 Discussion

All stimulation of the brain will result in the deposition of thermal energy, whether through the Joule heating of implanted electrodes or the Joule heating of the resistive brain as currents are induced to flow. Although such thermal effects are generally inseparable with electrical stimulation, magnetic stimulation offers the ability to separate magnetic induction of electrical current effects on neural activity from thermal effects. We controlled the temperature changes by controlling the power dissipation through micro-magnetic coils during stimulation. During coil stimulation, we observed transient suppression of neuronal spiking activity in layer V pyramidal cells upon onset of stimulation (warming) and transient hyperactivity upon offset of stimulation (cooling). We found these effects to be purely thermal, as the same result during AC stimulation was obtained during DC stimulation experiments where there were no changing magnetic fields present. Since to our knowledge, cortical mammalian pyramidal cells in the rat are not sensitive to static DC magnetic fields, and in the absence of detecting magnetic induction effects on the spike timing of neurons, the transient changes in spiking with small temperature increases and decreases observed in this study could be attributed to purely thermal effects.

Temperature affects nearly all biological processes, and our computational model remains relatively simple by comparison with the vast experimental literature on temperature. Nevertheless, we found that augmentation of the fast voltage dynamics of the Hodgkin-Huxley equations with the slower dynamics of compartmental ion fluxes (28), and including additional temperature sensitivity through *Q*_10_ rate factors, was indeed sufficient to capture the driving dynamics of warming and cooling on model spiking in a qualitatively similar manner to our experimental observations.

Nonlinear systems can undergo critical transitions – bifurcations or tipping points – where rather than small responses to small changes in parameters, one can observe abrupt, large-scale changes in dynamical behavior (32). Bifurcation analysis of our computational model uncovered a previously undescribed bifurcation that accounts for the transient suppression of spiking during warming, and the excitation during cooling. Furthermore, bifurcations in the steady state dynamics leading to stable firing suppression are described for higher temperatures than explored in these experiments, predicting suppression of spiking with the appearance of a stable state at these more elevated temperatures. These latter stable states will require future experimental exploration and validation.

The rate dependency of the bifurcations or tipping points observed as in Figure 4c and 4d share a remarkable similarity with other warming phenomena. In particular, there has been much interest in planet-scale transitions in response to crossing temperature thresholds by global warming (33). Nevertheless, it has been recently recognized that the rate of change of warming might be the critical determinant of an abrupt transition – so called rate-dependent tipping points in open (driven) systems (34).

In neuronal systems, excitability was a paradigm for tipping points with respect to the rate of change of an input; indeed, a gradual depolarization of the neuronal membrane might be accommodated without disruption of the resting state of the cell, but a larger rate of depolarization would lead to an excitable spike in voltage response (35). In climate change literature, there is described a compost-bomb instability, where beyond a certain critical rate of warming, the increase in soil carbon decomposition rate with temperature can no longer be dissipated fast enough, and a catastrophic release of peatland soil carbon ensues (36). Such rate-dependent excitability responses can be seen with not just stable equilibria but also with limit cycle dynamics. A slow enough increase in temperature can cause such systems to adiabatically follow the slowly drifting equilibria or limit cycle, but beyond a certain critical rate of warming, a tipping point with a large transient occurs.

In our system, summarized in Figure 4c, at slow rates of warming the neuron will change equilibrium points but maintain a stable limit cycle during the transition. Above a certain rate of change in temperature, there is a large excursion of variable values that veers away from, but slowly returns to the same eventual stable limit cycle reached with slow adiabatic warming. In Figure 4c, we see that rapid warming results in the electrogenic extrusion of *Na^+^* and internalization of *K*^+^, which cannot be dissipated on the same fast time scale – the cell loses its limit cycle and silencing occurs as the cell hyperpolarizes. The opposite occurs upon cooling. This thermal rate-dependent tipping point of the mammalian neuron shares a remarkable qualitative similarity with the warming instability of Siberian peatlands. The rates to induce an ecological tipping point are of the order of 0.1°C per year for 20 years, whereas in our thermal neuronal model the tipping point is observed for warming of 1°C for 5 seconds.

While our relatively simple temperature modified 2-compartment ion-conserving Hodgkin-Huxley type model did capture temperature effects on spiking, we did not attempt to include many additional factors such as temperature-induced changes in the cell membrane’s capacitance (37; 38; 39), nanoporation (39), changes in *Ca*^2+^ currents (40; 41; 42; 43) and the subsequent activation of calcium-dependent *K* (e.g., BK) channels (44), or G-protein coupled receptors (45). To focus on the single neuron effects from temperature, we did not account for the temperature dependency of the glial [*K*^+^]*_o_* uptake nor of extracellular ionic diffusion. Cationic transient receptor potential (TRP) channels (46) are expressed in mammalian cortical cells, modulated by temperature with *Q*_10_ values ranging from 5 to 100 (46), and can contribute to suppression of cellular spiking activity in response to heat pulses (44). Nevertheless, none of these complex unmodeled biological features were necessary to account for the transient thermal suppression and cooling hyperactivity seen in our experiments.

The Na-K ion pumps and the currents they generate are one of the dominant energy consuming elements of the brain (47). It therefore is logical that given their *Q*_10_ value that they might be an important component of thermal effects. Carpenter et al. (48), in their study on Aplysia neurons, demonstrated that blocking Na-K pump function with ouabain abolishes the hyperpolarization observed following warming, consistent with prior experimental results (2). Similarly, in our present work the effect of the Na-K pump dynamics was sufficient to account for the transient dynamics observed.

The energy neutral ion co-transporters are important in modulating excitability levels and response to temperature changes. The conversion from the immature balance of KCC2 versus NKCC1 to a mature balance accounts for the ‘GABA switch’ observed from excitation to inhibition induced by gamma-aminobutyric acid (GABA) effects on chloride channel current (49; 50; 51; 52; 53; 54). Because KCC2 extrudes *Cl*^−^, intracellular [*Cl*^−^]*_i_* levels decrease with development as expression of KCC2 increases, increasing the reversal potential *E_Cl_* (Fig. 5c), and pushing the cell into a more hyperpolarized state; our experiments were conducted in brain cells that were towards the end of this maturation period (55). Previous work has demonstrated heat-induced increases of neural activity in immature animals (56; 57). We demonstrated that lowering KCC2 levels in our model resulted in a switch from suppression to excitation as temperature is raised 1-2 degrees Celsius above the physiological baseline. It is intriguing to speculate whether such effects may shed light upon childhood febrile seizures, where seizures are often transient phenomena during the rising temperature phase of a febrile episode.

Although our single-cell experimental and modeling results provide a converging picture of how small temperature increases suppresses spiking temporarily in neuronal membranes, there are additional structural features of the experiments that could contribute to the findings observed. Experimentally, the coil element was lithographically deposited on the underside of the recording chamber, which created a temperature gradient across the thickness of the cortical section, and consequently across different neural populations within our coronal brain slices. A more complex analysis would consider the spatiotemporal profile (58) of the heat diffusion through the tissue.

The lack of detection of frequency differences, spike height changes, or spike phasic entrainment by oscillating magnetic fields is or course limited by the precision of our timing measurements. Given our high bandwidth of recording (100kHz, digitally filtered to 10kHz), any residual magnetic induction effect that we failed to detect would have to be extremely small. There is postulated to be a physical threshold limit of electric interaction with neurons – the so called ‘molecular shot noise’ of the neuronal membrane (59; 60; 61) – which our findings suggest we were within during these experiments.

A further advantage of *in vitro* brain slice preparations is that temperature can be controlled without the heat-sink effect and any blood flow response to temperature within the neuro-vascular unit. Such vascular interactions will render the thermal responses to local heating more complex, but will be essential to consider when modeling thermal effects of stimulation of the intact brain.

Because thermal effects are ubiquitously present when stimulating neural systems, it is important to ask what can help distinguish magnetic induction effects in experiments without thermal measurements and controls. We posit that orientation specificity of the neuronal response with respect to coil or electrode geometry should be present in magnetic or electrical stimulation. It is also possible to have increased heat transfer to neurons through the geometrical orientation of coil or electrodes on the scale of the neurons or networks being stimulated. Another factor in magnetic or electrical induction of neuronal responses is latency – electromagnetic radiation travels faster than heat propagates in neuronal tissue. Using a relatively large coil with respect to individual neuronal geometry, Pashut et al (19) demonstrated orientation specificity of neurons with respect to the coil geometry, and short latency responses (sub-millisecond), in addition to what appeared to be a refractory period (~150 ms) after single stimuli (at large field strengths, 0.4-0.9 Tesla). Using a multi-turn micro-coil, Bonmassar et al (16) found orientation specific short latency activation (~1 ms), followed by a more complex longer-lasting effect (subsequent suppression and activation over 250 ms following initial activation). Using single-loop micro-wires, Lee et al (18) demonstrated short latency (~1 ms) orientation specific activation of action potentials in cortical neurons. Yet there are examples in the literature of slower, more complex responses to magnetic stimulation that may at least partially reflect a thermal component. In Lee et al (62), prolonged sinusoidal stimulation suppressed activity at longer time scales in most subthalamic neurons studied with microcoil stimulation, with an interesting relationship to stimulation frequency and time to neuronal silencing. In Lee et al (63), there were observed short latency activation and suppression that were orientation specific, but additionally longer time scale (10s of seconds) neuronal activity changes that could also be dependent on whether a neuron had recently been stimulated. Also, since these varied reports in literature were observed from different sub-regions of brain, this raises the possibility that different neuronal sub-populations or cell-types may respond differently to thermal and magnetic stimulation.

The experimental and theoretical findings of this present work demonstrate that stimulation of the brain must take into account small thermal effects that are ubiquitously present in stimulation of the brain. More sophisticated models of electrical current interaction with neurons combined with thermal effects will lead to a better understanding of the effect of stimulation on neuronal response, and lead to more precise control of neuronal circuitry.

## 4 Materials and Methods

### 4.1 Experimental Design

We performed a series of experiments delivering thermal and magnetic stimulation to neocortical tissues with custom microscale electromagnetic coils. The experimental protocol was designed from 3-epoch trials alternating from DC current (thermal stimulation) to AC current (magnetic plus thermal stimulation) and reversed as shown in Fig. 2b (see also Fig. S1). This stimulation technique was implemented to control for the thermal energy delivered to the tissue via Joule heating of the micro-coil during the brief but high intensity current pulses required to induce magnetic and electric fields.

All experiments were performed in accordance with and approval from the Institutional Animal Care and Use Committee of The Pennsylvania State University. Neocortical slices were obtained from male Sprague-Dawley rats aged P11 to P14. Briefly, the animals were deeply anesthetized with diethyl-ether and decapitated, the brain removed and coronal slices were sectioned from occipital neocortex.

### 4.2 Slice preparation

After decapitation, the whole brain was quickly and carefully removed to chilled (4°C) artificial cerebrospinal fluid (ACSF) for 60 seconds containing the following (in mM): 126 NaCl, 2.5 KCl, 2.4 CaCl_2_, 10 MgCl_2_, 1.2 NaH_2_PO_4_, 18 NaHCO_3_, and 10 dextrose. ACSF was saturated with 95% O_2_ and 5% CO_2_ at room temperature for 1 hour before the dissection with an osmolality range from 295-310 mOsm and pH from 7.20-7.40. Coronal occipital cortical slices were cut with a vibratome on the rostro-caudal and medio-lateral coordinates of bregma −2 to −8 mm and lateral 1 to 6 mm, respectively. The first cut was made 1100 μm deep from the caudal surface and discarded, and 3 slices were taken from the brain each 300 μm thick. After cutting, slices were transferred to a chamber containing ACSF and allowed to recover for 30 minutes at 32-34° C, then incubated at room temperature (20-22° C) for an additional 30 minutes prior to recording. In order to examine the modulation of spiking in layer 5 pyramidal neurons, network spiking activity was supported by modified ACSF consisting of (in mM) 121.25 NaCl, 6.25 KCl, 1.5 CaCl_2_, 0.5 MgCl_2_, 1.25 NaH_2_PO_4_, 25 NaHCO_3_, and 25 dextrose, perfused over the slice at a rate of 2.8-3.0 mL/min at 30°C during recordings.

### 4.3 Microscale magnetic coil fabrication

The micro-coils (Fig. 1b) were fabricated on a 150 μm thick glass cover slip that formed the floor of our slice recording chamber. First the cover slip is cleaned with acetone, isopropyl alcohol and rinsed with deionized water followed by an O2 plasma ashing to remove organic residue. Next, a conductive seed layer of 20 nm thick Cr followed by 120 nm Au is evaporated for copper electroplating. To enable electroplating of thick copper coils a 28 μm thick MEGAPOSIT SPR 220 photoresist (Shipley Company, Marlborough, MA) is spun on the slides in two coats at 1500 RPM followed by a softbake. To minimize cracking and bubbling of the resist the softbake is conducted on two hotplates first at 95°C for 90s, then 115°C for 120s and finally back down to 95^°^C for 90s. The photoresist is exposed at a dose of 1200 mJ/cm^2^ on a SUSS MA/BA6 contact aligner (SÜSS MicroTec SE, Garching, Germany) in 15 cycles with 15s delays to reduce the risk of bubbling and cracking of the resist. This is followed by a minimum 30 min wait time before being developed in MICROPOSIT MF CD-26 (Shipley Company, Marlborough, MA) for 11 minutes. A final O2 plasma ashing is then performed to remove any remaining photoresist in the coil traces. The copper coil traces are then electroplated in a beaker using a high purity 99.99% copper anode and a Technic Copper FB Bath plating solution (Technic Inc., Cranston, RI) at 7 mA with a 20% duty cycle for 4 hours, followed by palladium electroplating to cap the copper with a non-oxidizing noble metal. The palladium is electroplated using Technic Pallaspeed RTU bathing solution (Technic Inc., Cranston, RI) using a Pt anode (Pd is dissolved in the solution) at 7 mA with a 20% duty cycle for 20 minutes. The photoresist is stripped using RemoverPG (MicroChem, Newton, MA) and the thin Cr/Au seed layer is etched away using an Ar/SF6 reactive ion etch. Finally an electroless Au is deposited using a gold potassium cyanide solution at 90°C for 2 hours. The coils are measured to be between 23 μm and 26 μm thick. This type of non-uniformity is typical of a beaker electroplating set-up. The resistance of the micro-coils used were measured between 0.43 - 0.60 Ω with a multimeter (Keithley 2000, Keithley Instruments, Solon, OH) before and after each experimental trial.

### 4.4 Electrophysiology, magnetic field and thermal application

The recording chamber (RC-22C; Warner Instruments, Holliston, MA) was fitted with a 150 μm thick cover slip and the chamber floor modified with lithographically deposited micro coils on the bottom (dry) side of the glass as shown in Figure 1a,b. Patch pipettes (4-6 MΩ) were pulled from thick-walled borosilicate glass capillaries with filaments (OD=1.5 mm, ID=0.86 mm; Sutter Instruments, Novato, CA) and filled with perfusate. Layer V pyramidal cells were targeted under visual control (Fig. 1a) and the patch pipette was adhered to the soma of the cell in loose patch configuration (25) (5-10 MΩ seal) in order to minimize the possibility of shunt current entering the cell from the pipette during magnetic stimulation. To drive magnetic stimulation, a function generator (Keithley 3390; Keithley Instruments, Solon, OH) supplied the AC input signal to an audio amplifier (PB717X, gain 2.16X; Pyramid Inc., or AE Techron 7224, gain 1.43X; AE Techron, Elkhart, IN, for high frequency stimulation up to 200kHz) which then amplified the signal sent to the coil. For thermal-only stimulation a DC power supply (Keithley 4590; Keithley Instruments, Solon, OH) was connected to the micro coil to supply heat to the tissue (stimulation details found in Table 2). Neuronal spiking was recorded (MultiClamp 700A; Axon Instruments, San Jose, CA) under perfusion with the modified ACSF and only actively spiking cells were tested with experimental stimulation. A Labview interface was used to low-pass filter and amplify the thermal signature recorded both at the surface of the chamber floor glass (wetted side) and nearby (~50 μm) from the patched cell using multiple highly sensitive (68 μV/°C, diameter 76 μm) micro E-type thermocouples (5SRTC-TT-E-40-36, Omega Engineering Inc.) as shown in Fig. 1a.

**Table 2:**
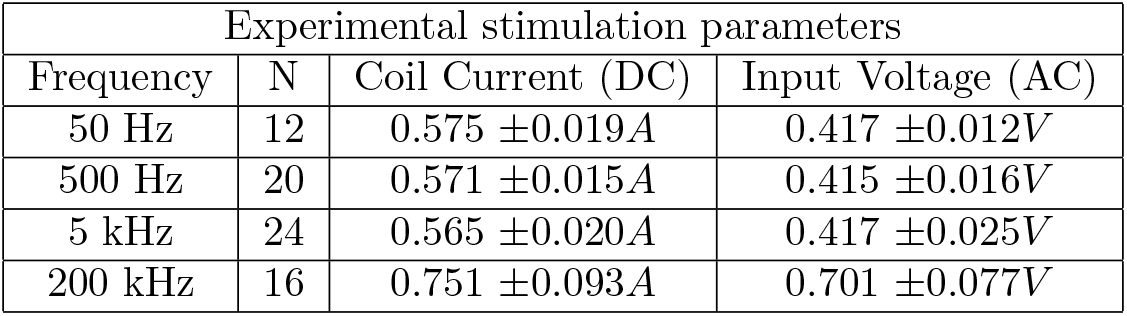
Summary of stimulation currents used during each frequency tested. The micro-coils used during stimulation ranged from 0.43-0.60 Ω.

Fig. S1 shows a typical experimental result. During the blue colored blocks A and E, no current was driven through the micro-coils, resulting in no additional heat or electromagnetic fields. In the green and red blocks B though D, the micro-coil is being driven in either AC or DC, producing heat and electromagnetic fields. The voltage placed across the micro-coil is shown in the top panel of Fig. S1a, and the temperature measured at the probe near the patched cell is shown in the middle panel in Fig S1a.

### 4.5 Statistical Analysis of Spike Entrainment

Spike entrainment refers to the phenomenon where spikes preferentially fire at certain phases of the applied AC sinusoidal stimulation. We used the Rayleigh Z test as the primary statistical tool to detect spike entrainment. For a set of *n* phase measurements, the Rayliegh Z metric is calculated by:

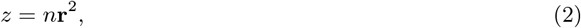

where **r** is a 2 dimensional vector with X and Y components,

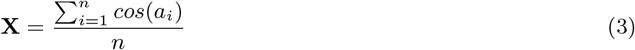

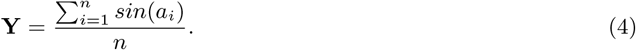

The null hypothesis of Rayleigh Z testing is that there is no preferred phase. The p-value can be approximated as exp 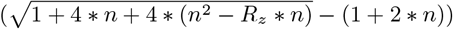, where *R_z_* is the Rayleigh Z statistic, and n is the number of measurements (64). We can also calculate Rayliegh Z for the DC blocks, using pseudophases. Pseudophases are measured by overlaying an AC block sinusoid on the DC spike data, and measuring the spike phases. These pseudophases serve as a control for the phases measured in the AC blocks.

First, we tested whether any AC blocks showed a preferential phase. Using the Benjamini-Hochberg procedure to control the false discovery rate at 10%, we found no blocks had statistically significant preferred phases.

We also accounted for the possibility that spike phase entrainment exists for short periods of time. For each block of phases and pseudophases, 99 surrogate spike data were generated by randomly shuffling the inter-spike intervals. For the ensemble of real and surrogate data, the time dependent power densities at the stimulation frequency in a moving window of 3 seconds were calculated using the Chronux package (65) (http://chronux.org). The confidence limit was set at 95th percentile of the ensemble, and we noted the temporal locations in each AC block where the power density of the real data passed the confidence limit. We grouped the time windows at these temporal locations with others if any part of them overlapped. We then obtained groups of p-values from Rayleigh-Z tests performed on the resulting combined candidate time windows, and again controlled for false discovery rate by applying the Benjamini–Hochberg method separately for phases and pseudophases at each stimulation frequency. If this returned any true-phase candidates as potentially significant at a particular stimulation frequency, we compared the distribution of the Rayliegh-Z p-values calculated from phases to those calculated from the pseudophases at that frequency in two ways: First, the Anderson-Darling test was used to test if the two samples come from different distributions. Second, a one-tailed Wilcoxon rank-sum test was used to see if the two distribution have different medians. Both tests failed to show greater degree of spike entrainment in AC blocks compared to DC blocks. Results are summarized in Table 1.

### 4.6 Full Model

Our model is based on the unification model from Wei et al 2014 (28): a single compartment spherical neuron model that expands the Hodgkin-Huxley model (29) to account for relevant structural micro-anatomy, conservation of charge and mass, energy balance, and volume changes. It consists of a single compartment spherical neuron with a thin extracellular space that is connected to the bath *in vitro* (or to a capillary *in vivo*) through a diffusion channel.

The membrane potential, *V*, is defined by:

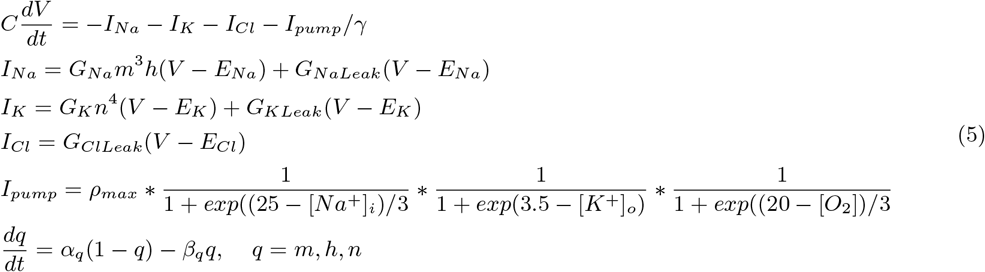

where *I_Na_, I_K_*, and *I_cl_* are the ohmic sodium, potassium, and chloride currents, respectively. *I_Na_* and *I_K_* consist of voltage-gated and leak currents, while *I_Cl_* consists solely of leak current. *G_Na_* and *G_K_* are maximal *Na*^+^ and *K*^+^ voltage-gated conductances, and *G_NaLeak_*, *G_KLeak_*, and *G_ClLeak_* are *Na*^+^, *K*^+^, and *Cl*^−^ leak conductances, respectively. We use conversion factor *γ* = *S*/(*F_v_i__*) to convert *I_pump_* from [mM/s] to [μA/cm^2^], where *S* is the surface area of the cell, *F* is the Faraday constant, and *v_i_* is the intracellular volume of the cell.

Reversal potentials are calculated with Nernst equations:

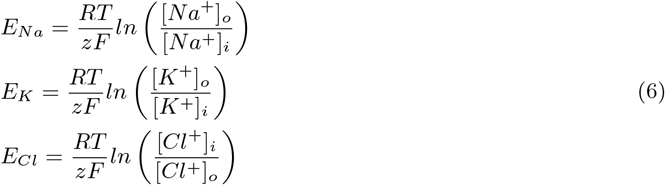

The KCC2 cotransporter was modeled as:

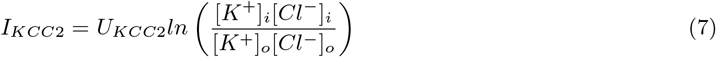

Values and descriptions of the parameters are given in Table 3, and the full description of the model is given in (Supplementary Material).

**Table 3:** Values of parameters used in the model.

| Parameter | Base Value | Description |
| --- | --- | --- |
| T | 30°C | Temperature |
| C | 1μF/cm<sup>2</sup> | Membrane capacitance |
| G<sub>Na</sub> | 45mS/cm<sup>2</sup> | Maximal conductance of sodium current |
| G<sub>K</sub> | 25mS/cm | Maximal conductance of potassium current |
| G<sub>NaLeak</sub> | 0.0247mS/cm<sup>2</sup> | Conductance of leak sodium current |
| G<sub>KLeak</sub> | 0.05mS/cm<sup>2</sup> | Conductance of leak potassium current |
| G<sub>ClLeak</sub> | 0.1mS/cm<sup>2</sup> | Conductance of leak chloride current |
| β<sub>0</sub> | 7 | Ratio of the initial intra-/extracellular volume |
| ρ<sub>max</sub> | 0.8mM/s | Maximal Na/K pump rate |
| G<sub>glia,max</sub> | 5mM/s | Maximal glial uptake strength of potassium |
| ε<sub>K,max</sub> | 0.25 s<sup>-1</sup> | Maximal potassium diffusion rate |
| ε<sub>Na,max</sub> | 0.1705 s<sup>-1</sup> | Maximal sodium diffusion rate |
| ε<sub>Cl,max</sub> | 0.2597 s<sup>-1</sup> | Maximal chloride diffusion rate |
| ε<sub>O</sub> | 0.17 s<sup>-1</sup> | Oxygen diffusion rate |
| K<sup>+</sup><sub>bath</sub> | 6.25mM | Normal bath potassium concentration |
| Na<sup>+</sup><sub>bath</sub> | 147.4mM | Normal bath sodium concentration |
| Cl<sup>-</sup><sub>bath</sub> | 131.4mM | Normal bath chloride concentration |
| α | 5.3g/mol | Conversion factor |
| O<sub>2bath</sub> | 32 mg/L | Normal bath oxygen concentration |
| U<sub>Kcc2</sub> | 0.3mM/s | Maximal KCC2 cotransporter strength |
| U<sub>Nkcc1</sub> | 0.1mM/s | Maximal NKCC1 cotransporter strength |

### 4.7 Temperature modeling

The main premise of our temperature model is that cellular processes will accelerate when a moderate amount of heat is introduced. We used *Q*_10_ factors to model these accelerated rates. The *Q*_10_ factor of a process describes the change of the reaction rate when the temperature is changed by 10^°^C. For an arbitrary temperature difference, the *Q*_10_ is calculated as:

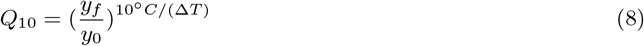

where Δ*T* is the temperature change and *y* is the rate of a process. With a known *Q*_10_, the rate at a new temperature *T* is given by:

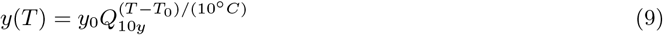

Heat propagation is not instantaneous, and we assume that the temperature exponentially decays, given by:

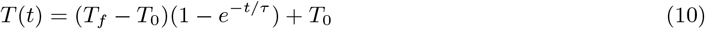

where *τ* is the time constant of the exponential decay. Combining equations 9 and 10 we get the time dependent rate equation:

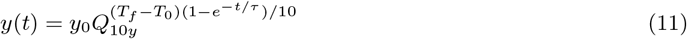

Using typical values reported in past literature (Table 4), We estimated *Q*_10_ model parameter values as in (Table 5). All full model computations were done in MATLAB (Mathworks).

**Table 4:** List of *Q*_10_’s reported in literature

| Subject | Parameter | Q<sub>10</sub> |
| --- | --- | --- |
| Giant squid axon(3) | Membrane conductivity | 1.3 |
|  | Rate of change of gating variable | 3 |
| Rabbit sciatic nerve(66) | Time constant for h | 1/3 |
|  | Conductivity of Na channels | 1.7 |
| Rat omohyoid muscle(67) | Time constant for n | 1/2 |
| Guinea pig hippocampal CA1 pyramidal neurons(68) | Inverse input resistance | 1.72 |
| Rat brain(69) | Immature NaKATPase activity | 2.58 |
|  | Adult NaKATPase activity | 3.05 |
| Myelinated nerve fibres of *Xenopus laevis*(70) | $\alpha_m, \beta m$ | 1.8, 1.7 |
| | $\alpha_h, \beta h$ | 2.8, 2.9 |
| | $\alpha_n, \beta n$ | 3.2, 2.8 |
|  | Sodium channel conductance | 1.3 |
|  | Potassium channel conductance | 1.2 |

**Table 5:** *Q*_10_’s used in the base model

| Parameter | $Q_{10}$ |
| --- | --- |
| $\frac{dm}{dt}$ | 1.75 |
| $\frac{dh}{dt}$ | 2.85 |
| $\frac{dn}{dt}$ | 3 |
| $\rho_{\max}$ | 3 |
| $G_{Na}$ | 1.7 |
| $G_K$ | 1.7 |
| $G_{NaLeak}$ | 1 |
| $G_{KLeak}$ | 1 |
| $G_{ClLeak}$ | 1 |
| $U_{Nkcc1}$ | 3 |
| $U_{Kcc2}$ | 3 |

### 4.8 Reduced Model Bifurcation Analysis

Bifurcations of the system’s long-term behaviors were studied using numerical continuation software. We used MatCont (71), an open-source solver built on MATLAB. In order to improve reliability of this analysis, we used a reduced version of the model (28) where we replaced three ion concentrations with functions of other concentrations, as shown in equations 12. In addition, we also confined the cell volume to be constant.

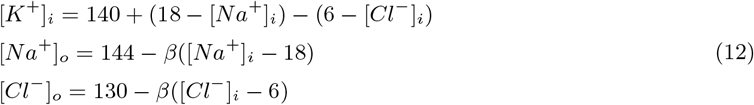

## 5 Acknowledgements

We are greateful to M. M. Norton for his helpful discussions. Supported by NIH grants 1R01EB01464, 1R21EY026438 and 2R01EB014641.

## Supplementary Information

### 6.1 Details of the Unification Model

Model parameteters are given in Table 3.

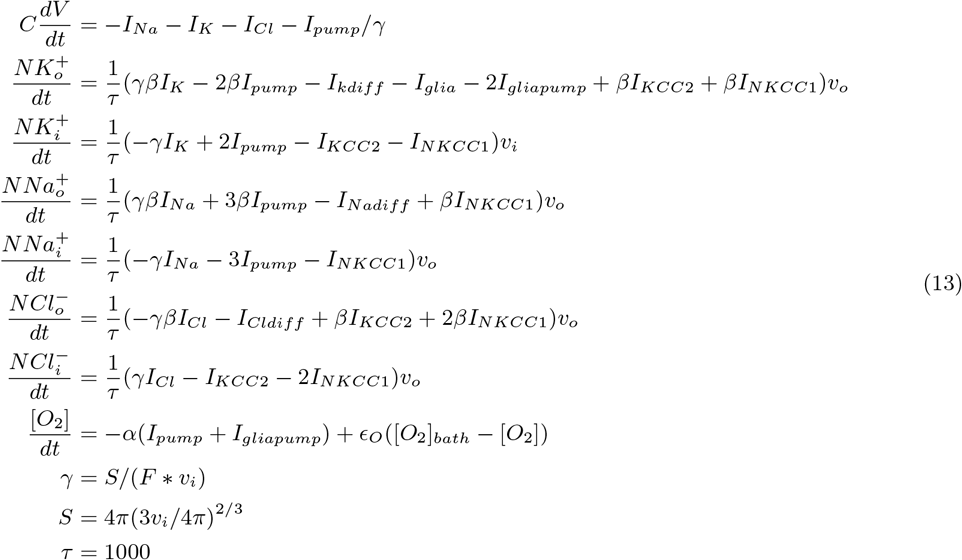

*τ* is the conversion factor from seconds to milliseconds, *β* is the ratio of intracellular volume to extracellular volume, S is the surface area of the cell, *F* is the Faraday constant, and *γ* is the conversion factor to convert concentration change (mM/s) into current density (μA/cm^2^).

The ionic currents and Nernst potentials are:

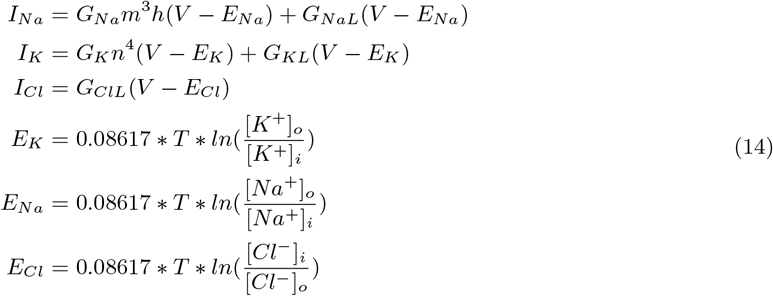

where *T* is the temperature in K.

Rate of change of gating variables are given by:

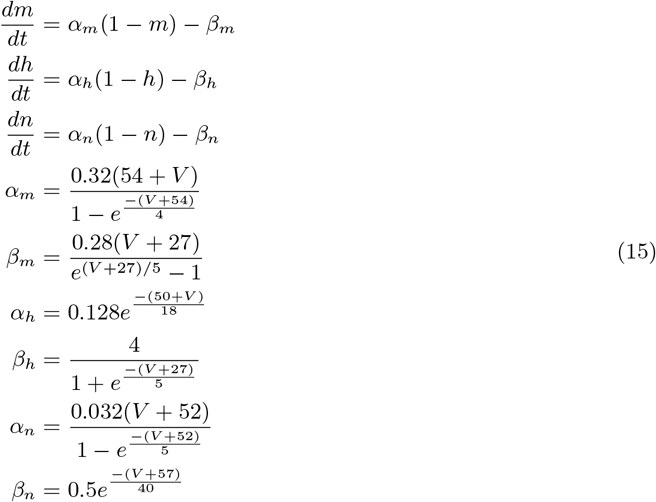

The pump currents and the glial currents are:

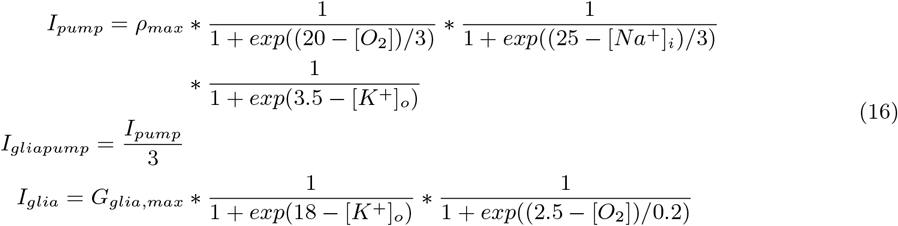

Cell volume:

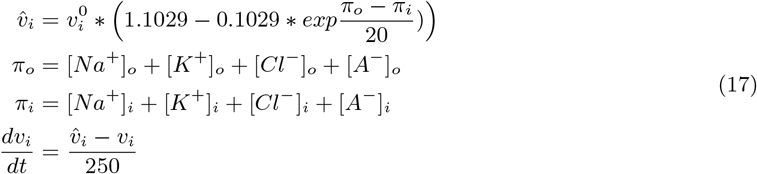

where [*A*^−^]*_i_* and [*A*^−^]*_o_* are the concentrations of impermeable anions in the intracelluar and extracelluar space, respectively. [*A*^−^]*_o_* is assumed to be 18 mM while [*A*^−^]*_i_* is calculated at the initial condition such the osmotic pressure is at an equilibrium.

The non-electrogenic cotransporters NKCC1 and KCC2 are given by:

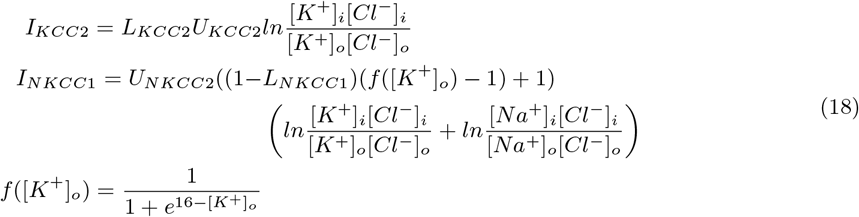

where *L*_*NKCC*_1__ and *L*_*KCC*_2__ are cotransporter levels. Diffusion to the extracullar space is given by:

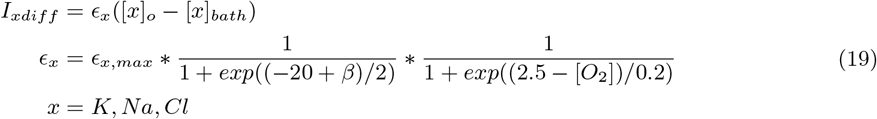

The model was initialized by running the model from the initial conditions as shown in Table S1 until the system reaches a steady-state.

**Table S1:**
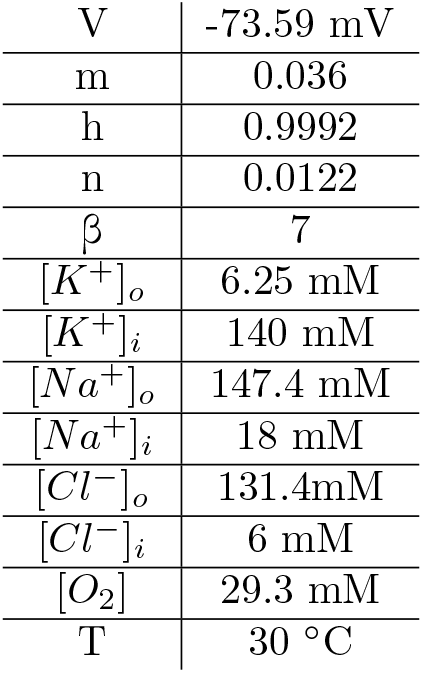
Coordinates used to initialize the model

|  |  |
| --- | --- |
| V | -73.59 mV |
| m | 0.036 |
| h | 0.9992 |
| n | 0.0122 |
| $\beta$ | 7 |
| $[K^+]_o$ | 6.25 mM |
| $[K^+]_i$ | 140 mM |
| $[Na^+]_o$ | 147.4 mM |
| $[Na^+]_i$ | 18 mM |
| $[Cl^-]_o$ | 131.4mM |
| $[Cl^-]_i$ | 6 mM |
| $[O_2]$ | 29.3 mM |
| T | 30 °C |

### 6.2 Stability of Equilibria in the 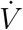 Curve

An extended version of the 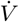 curve at the onset of the temperature step (Fig. 3**i**) is shown in Fig. S7. There exists three zeros at P1, P2, and P3 as shown, which are the equilibria of the system. Conventional wisdom tells us that an equilibrium is stable if the slope of the first derivative is negative. We can see that P1 and P3 have negative slopes and P2 has positive slope. While we can concude that P2 is an unstable equilibrium, this does not tell the full story for P1 and P3.

A true stable equilibrium will be stable against small perturbations to the gating variables. We tested the stability for small shifts in potential from *V_eq_* to *V* = *V_eq_* + *δV*. The stability of the gating variables at the equilibria was determined by calculating the eigenvalues of the Jacobians under these assumptions. For stable equilibria, the eigenvalues must all have negative real parts. These calculations were done in Mathematica (Wolfram Research, Inc. Champaign, Illinois).

The system of equations for the voltage and the gating variables is the same as the full model’s:

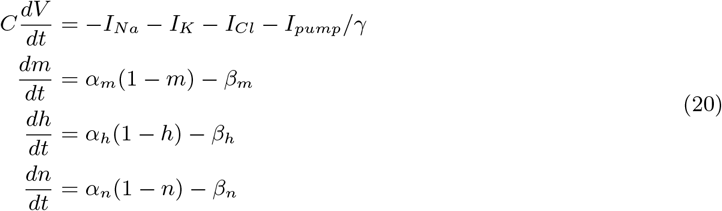

Near some *V* = *V_i_*, the Jacobian is calculated as the matrix:

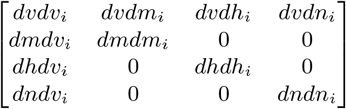

where the elements are given by:

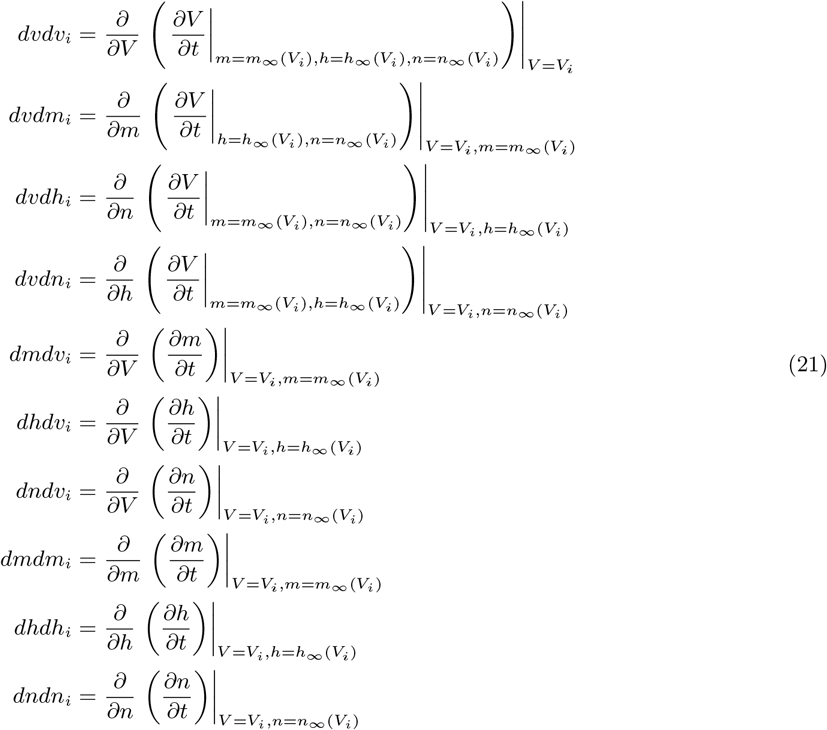

For point 1, all eigenvalues were real and negative, making this equilibrium stable. Point 3 also had all real eigenvalues, but only 2 of them were negative, thus making it unstable to small changes in the gating variables.

Analysis of the field map around these points confirms that point 1 is the only true stable point. Near these equilibria, *dm/dt* >> *dn/dt* >> *dh/dt* (Figure S8). Thus, we assume that for each gating variable, faster gating variables are at their steady-state values for the corresponding voltage (*x*_∞_(*V*)), and slower gating variables are at their steady-state values for the equilibrium voltage (*x*_∞_(*V_eq_*)).

## 6.3 Supplementary Figures

**Figure S1:**
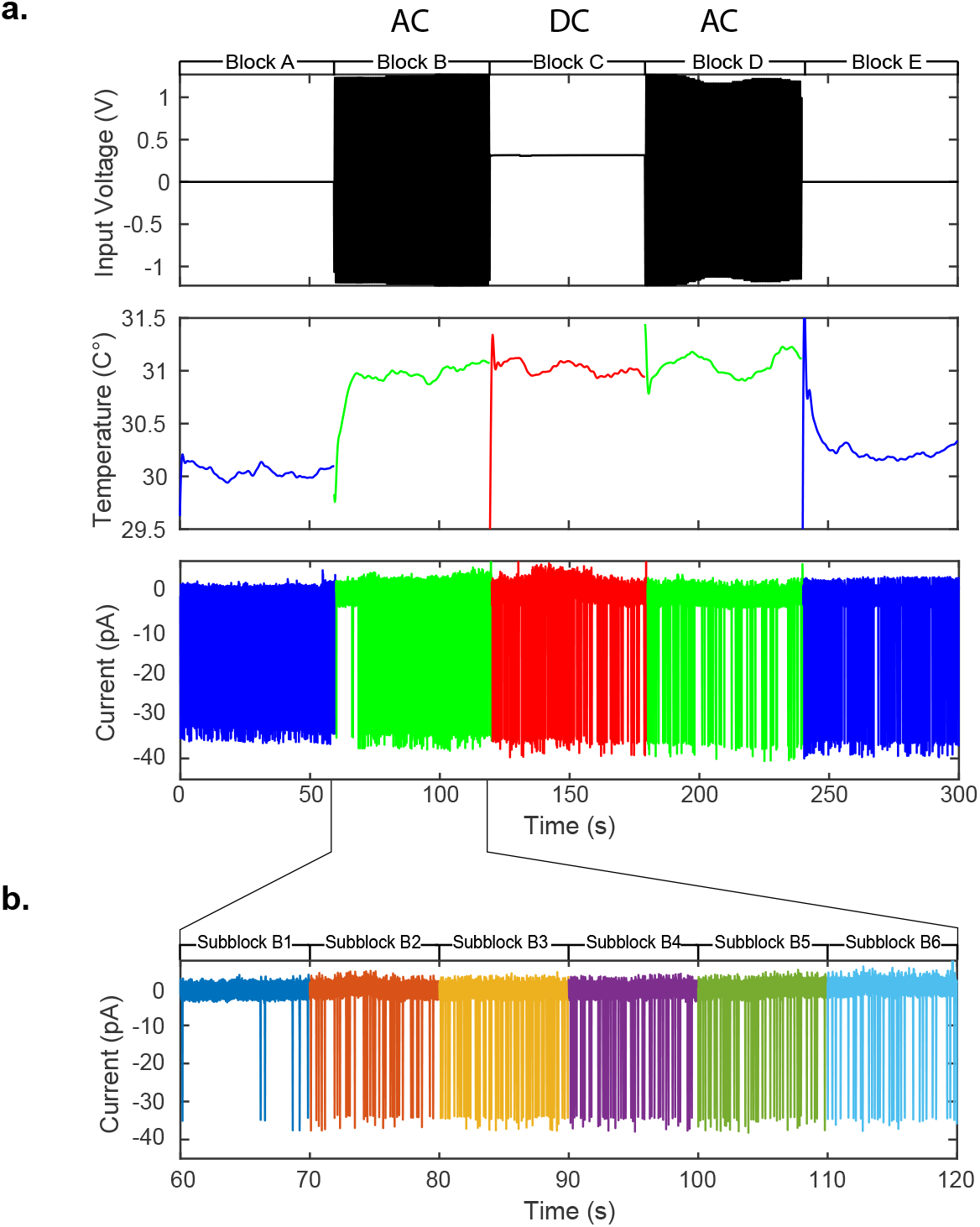
**a**, Representative traces of the stimulation voltage, temperature near the patched cell, and the patched cell recording from a block design of alternating AC and DC current stimulation to control for temperature changes. The recordings are broken into 5 temporal blocks, from A to E. A and E blocks are control blocks with no stimulation, blocks B, C, and D are stimulation blocks. B and D blocks always have the same stimulation paradigm (either AC or DC), while C blocks were stimulated in the other paradigm. **b** Each block was also further divided into sub-blocks of 10 seconds for subsequent statistical analysis.

**Figure S2:**
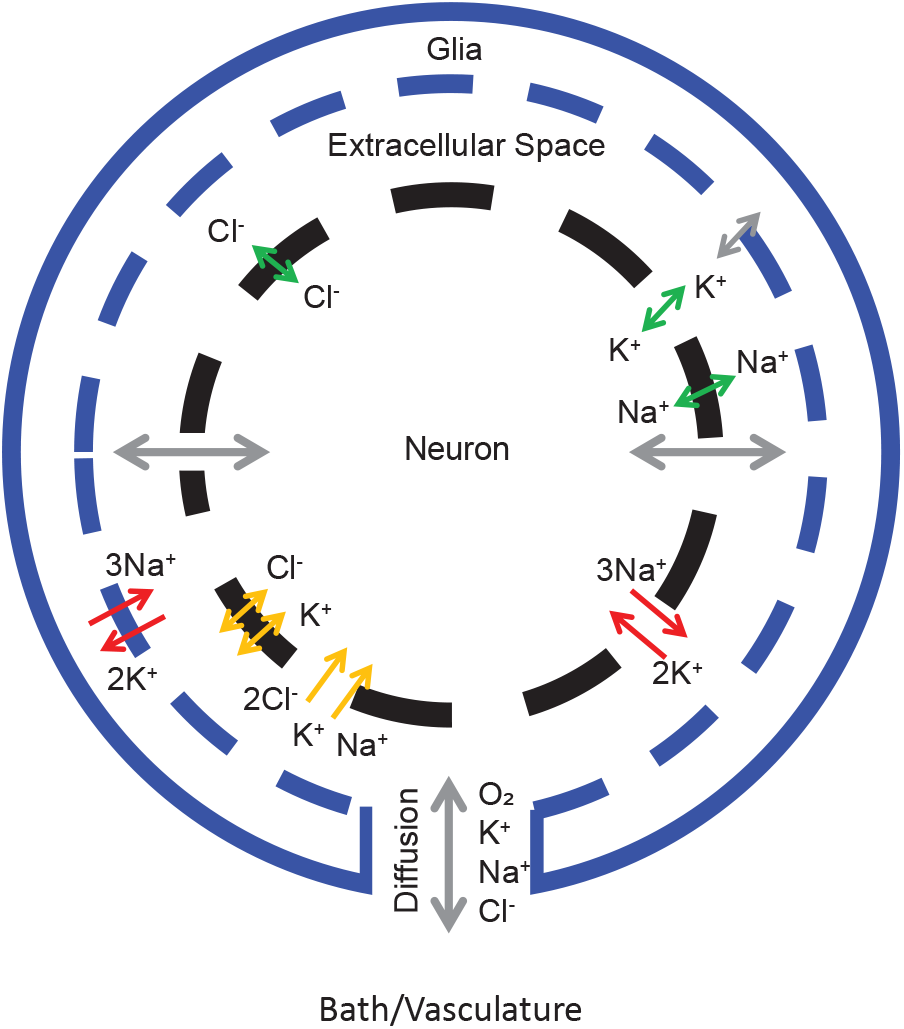
Model schematic. The model consists of a single compartment neuron that is separated from the bath by a thin extracellular space and glia. In addition to the Hodgkin-Huxley style ohmic currents (green arrows), the model includes volume changes, active pumps (red arrows) and cotransporters (yellow arrows). Gray arrows represent components not modeled to change with temperature, while colored arrows are temperature dependent.

**Figure S3:**
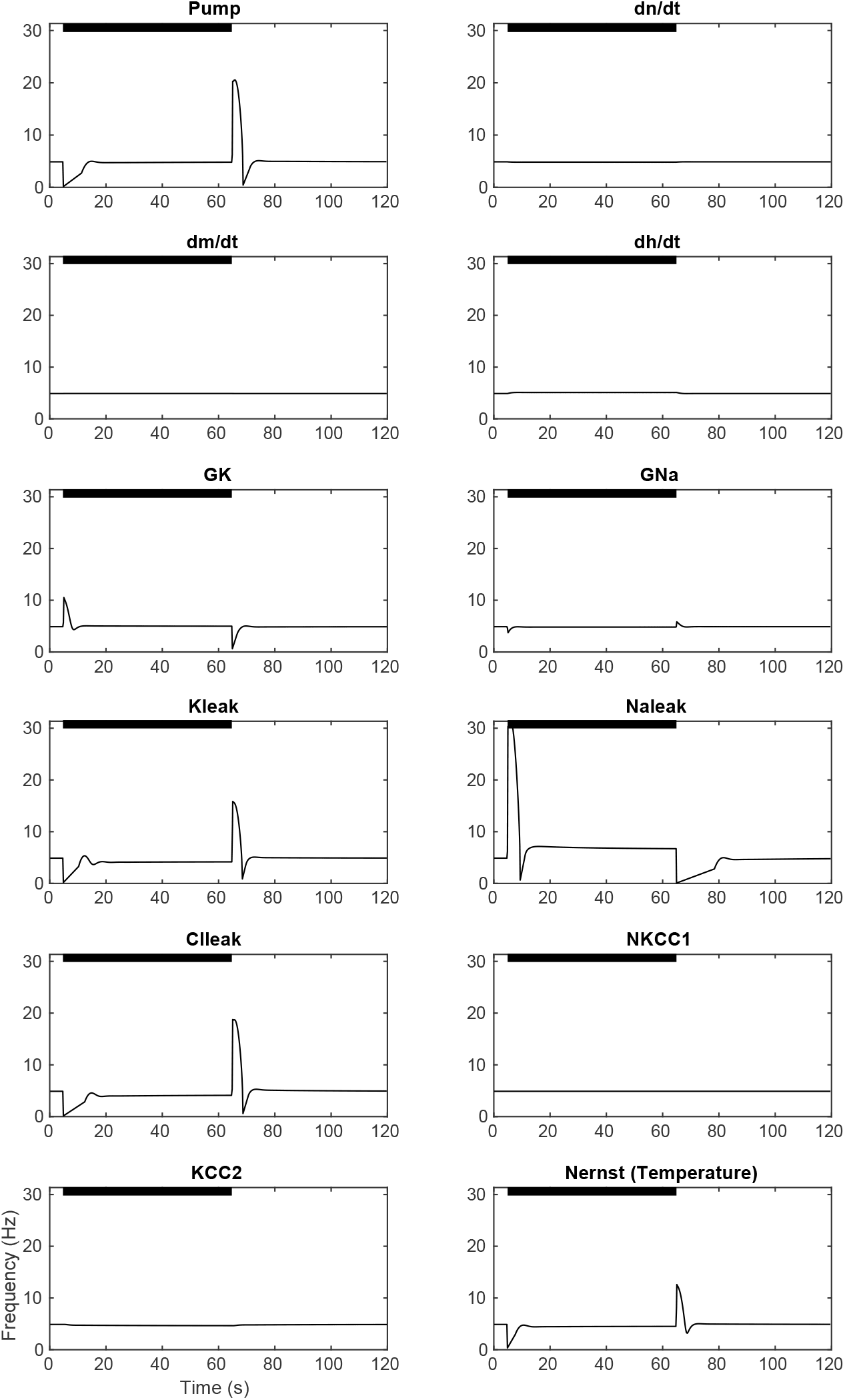
Frequency displayed by the model when each membrane parameter is individually increased by 10%. In the lower right, the Nernst potential temperature is increased by 1°C. The black bars indicate the times when the parameters are raised over their base values.

**Figure S4:**
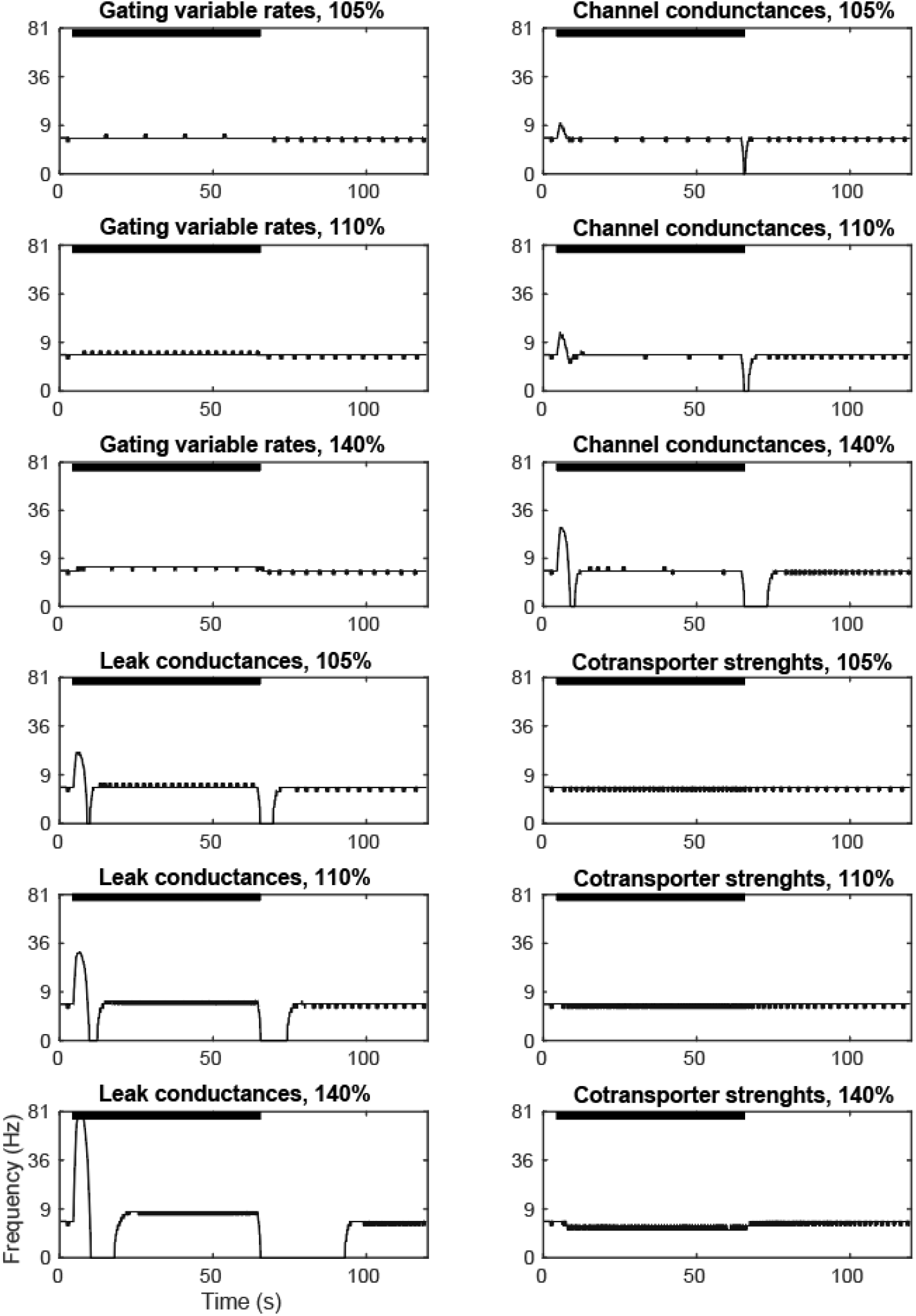
Frequency displayed by the model when groups of parameters are changed by the same amount to reflect similar *Q*_10_ changes. The black bars indicate the times when the parameters are raised over their baseline values.

**Figure S5:**
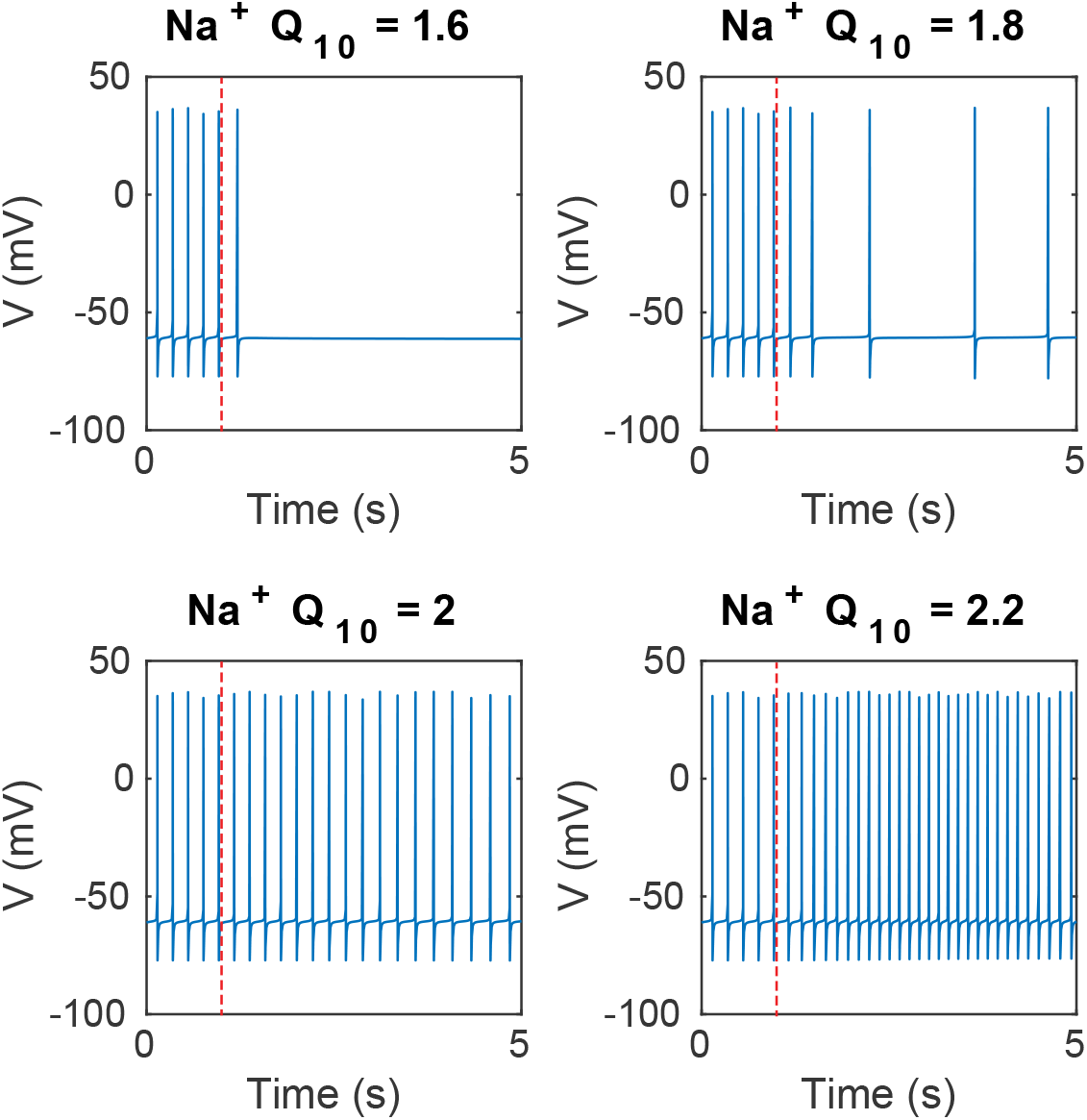
Model behavior when changing the temperature at 1 second (red dashed line) at various 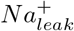 *Q*_10_ levels. 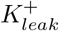 and 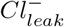 *Q*_10_’s are set at 3 (near their physiological observed limits, see Table 3). The model shows that 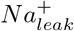 can counteract the silencing of 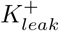 and 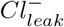 *Q*_10_, for values of 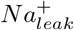 *Q*_10_ above 2/3 the value of the 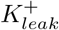 and 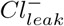 *Q*_10_. Because such a large deviation between *Q*_10_ values for similar processes are unlikely, leak is unlikely to be the driving force of the transient silencing observed in the experiments.

**Table S2:** Result of MANOVA for each 10 second sub-blocks. To account for any time dependent effects, we applied MANOVA to each sub-block separately. The independent variable is the stimulation frequency (0, 50, 500, 5000, and 200000 Hz), and the dependent variables are the normalized spike heights and density. Using Benjamini–Hochberg (72) false discovery analysis, we found that there are there were no statistically significant changes to the spike height or spike frequency at any driving frequency.

| Sub-block | Statistic | Value | F | df1 | df2 | p val | $\eta^2$ |
| --- | --- | --- | --- | --- | --- | --- | --- |
| B1 | Pillai's Trace | .091 | 1.081 | 8.000 | 182.000 | .379 | .045 |
|  | Wilks' Lambda | .911 | 1.079b | 8.000 | 180.000 | .380 | .046 |
| B2 | Pillai's Trace | .082 | .970 | 8.000 | 182.000 | .461 | .041 |
|  | Wilks' Lambda | .919 | .971b | 8.000 | 180.000 | .460 | .041 |
| B3 | Pillai's Trace | .091 | 1.079 | 8.000 | 182.000 | .380 | .045 |
|  | Wilks' Lambda | .910 | 1.091b | 8.000 | 180.000 | .372 | .046 |
| B4 | Pillai's Trace | .090 | 1.072 | 8.000 | 182.000 | .384 | .045 |
|  | Wilks' Lambda | .912 | 1.062b | 8.000 | 180.000 | .392 | .045 |
| B5 | Pillai's Trace | .049 | .570 | 8.000 | 182.000 | .802 | .024 |
|  | Wilks' Lambda | .952 | .566b | 8.000 | 180.000 | .805 | .025 |
| B6 | Pillai's Trace | .167 | 2.072 | 8.000 | 182.000 | .041 | .083 |
|  | Wilks' Lambda | .835 | 2.128b | 8.000 | 180.000 | .035 | .086 |
| C1 | Pillai's Trace | .030 | .344 | 8.000 | 182.000 | .948 | .015 |
|  | Wilks' Lambda | .970 | .341b | 8.000 | 180.000 | .949 | .015 |
| C2 | Pillai's Trace | .031 | .363 | 8.000 | 182.000 | .939 | .016 |
|  | Wilks' Lambda | .969 | .362b | 8.000 | 180.000 | .939 | .016 |
| C3 | Pillai's Trace | .026 | .303 | 8.000 | 182.000 | .964 | .013 |
|  | Wilks' Lambda | .974 | .300b | 8.000 | 180.000 | .965 | .013 |
| C4 | Pillai's Trace | .076 | .661 | 8.000 | 134.000 | .725 | .038 |
|  | Wilks' Lambda | .925 | .655b | 8.000 | 132.000 | .730 | .038 |
| C5 | Pillai's Trace | .076 | .662 | 8.000 | 134.000 | .724 | .038 |
|  | Wilks' Lambda | .925 | .656b | 8.000 | 132.000 | .729 | .038 |
| C6 | Pillai's Trace | .122 | 1.084 | 8.000 | 134.000 | .378 | .061 |
|  | Wilks' Lambda | .881 | 1.076b | 8.000 | 132.000 | .384 | .061 |
| D1 | Pillai's Trace | .150 | 1.355 | 8.000 | 134.000 | .222 | .075 |
|  | Wilks' Lambda | .855 | 1.346b | 8.000 | 132.000 | .227 | .075 |
| D2 | Pillai's Trace | .149 | 1.351 | 8.000 | 134.000 | .224 | .075 |
|  | Wilks' Lambda | .855 | 1.340b | 8.000 | 132.000 | .229 | .075 |
| D3 | Pillai's Trace | .123 | 1.094 | 8.000 | 134.000 | .371 | .061 |
|  | Wilks' Lambda | .881 | 1.083b | 8.000 | 132.000 | .379 | .062 |
| D4 | Pillai's Trace | .203 | 1.889 | 8.000 | 134.000 | .067 | .101 |
|  | Wilks' Lambda | .806 | 1.876b | 8.000 | 132.000 | .069 | .102 |
| D5 | Pillai's Trace | .145 | 1.312 | 8.000 | 134.000 | .243 | .073 |
|  | Wilks' Lambda | .860 | 1.295b | 8.000 | 132.000 | .252 | .073 |
| D6 | Pillai's Trace | .081 | .706 | 8.000 | 134.000 | .685 | .040 |
|  | Wilks' Lambda | .919 | .708b | 8.000 | 132.000 | .684 | .041 |

**Figure S6:**
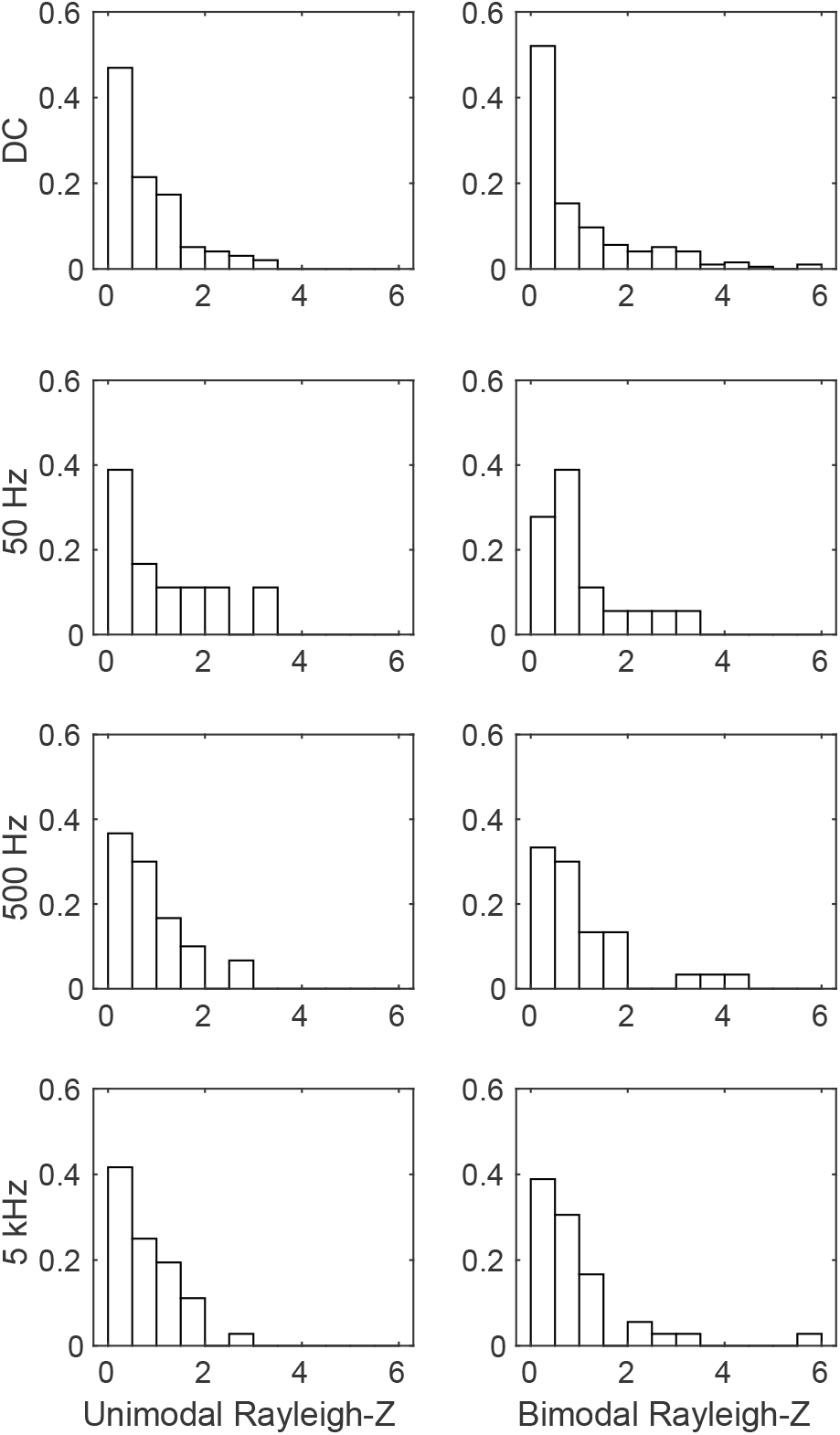
Histogram of Rayleigh-Z values from experiments, grouped by the stimulation frequency and the modality used in calculating the Rayleigh-Z (unimodal or bimodal). For DC and control blocks, the sinusoidal signal from the AC block was analytically extended to create a pseudophase for spike timing necessary for comparisons between all blocks. The distributions of Rayleigh-Z grouped by stimulation paradigm were then compared to each other using Kruskal-Wallis test, under assumptions of both uni- and bi-modality. No AC blocks showed statistically significant directionality compared to the DC blocks (p = 0.3887 for unimodal, p = 0.1721 for bimodal).

**Figure S7:**
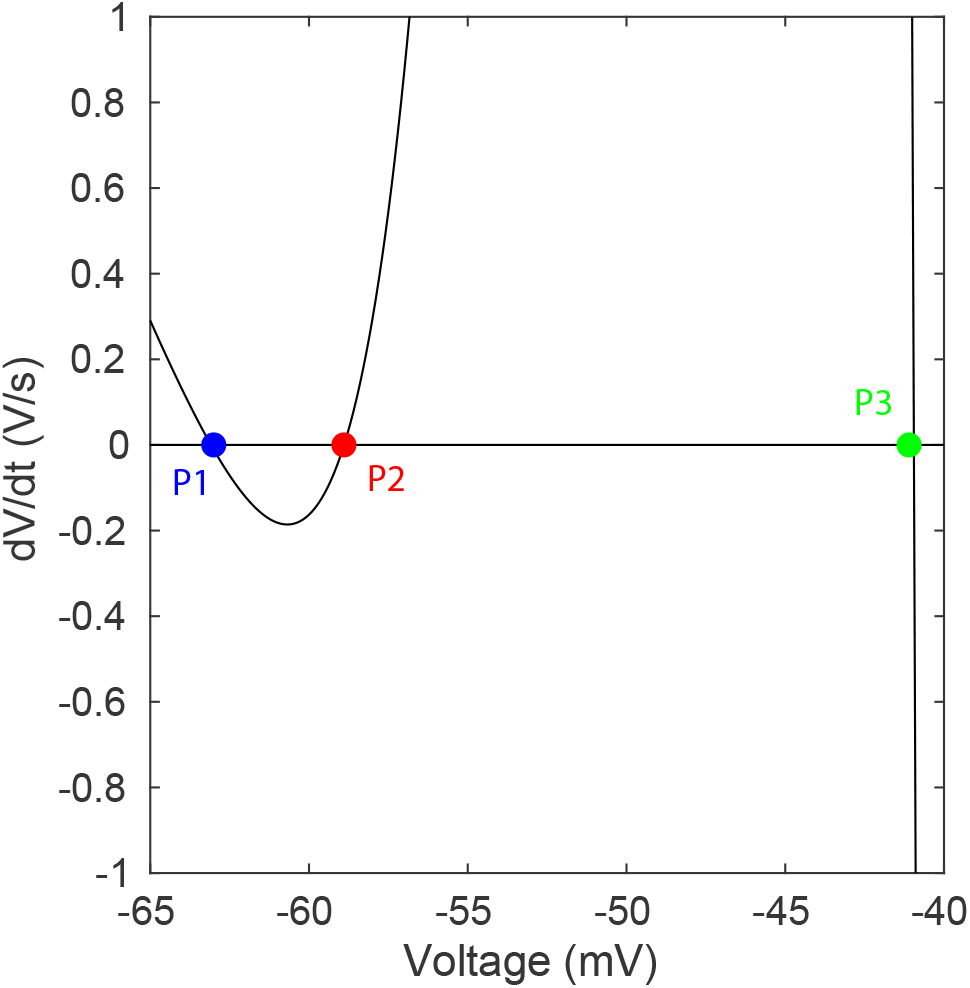
Locations of all three equilibria of the 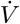 curve. P1 is a stable equilibrium, while P2 and P3 are unstable equilibria.

**Figure S8:**
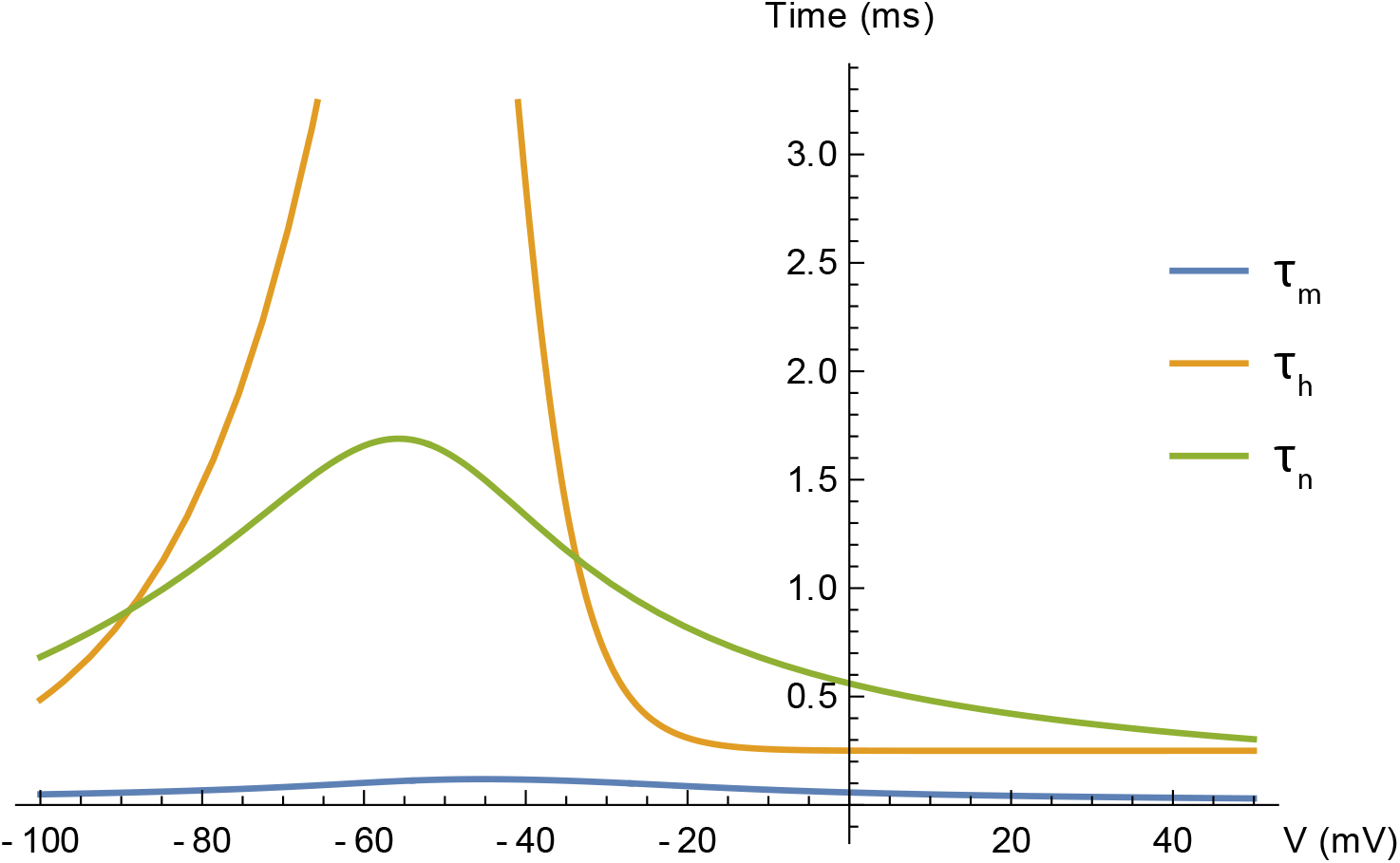
Time constants of the three gating variables m, n, and h as functions of membrane potential. Near the region of resting membrane potential, we can see that *τ_h_* >> *τ_n_* >> *τ_m_*.

**Figure S9:**
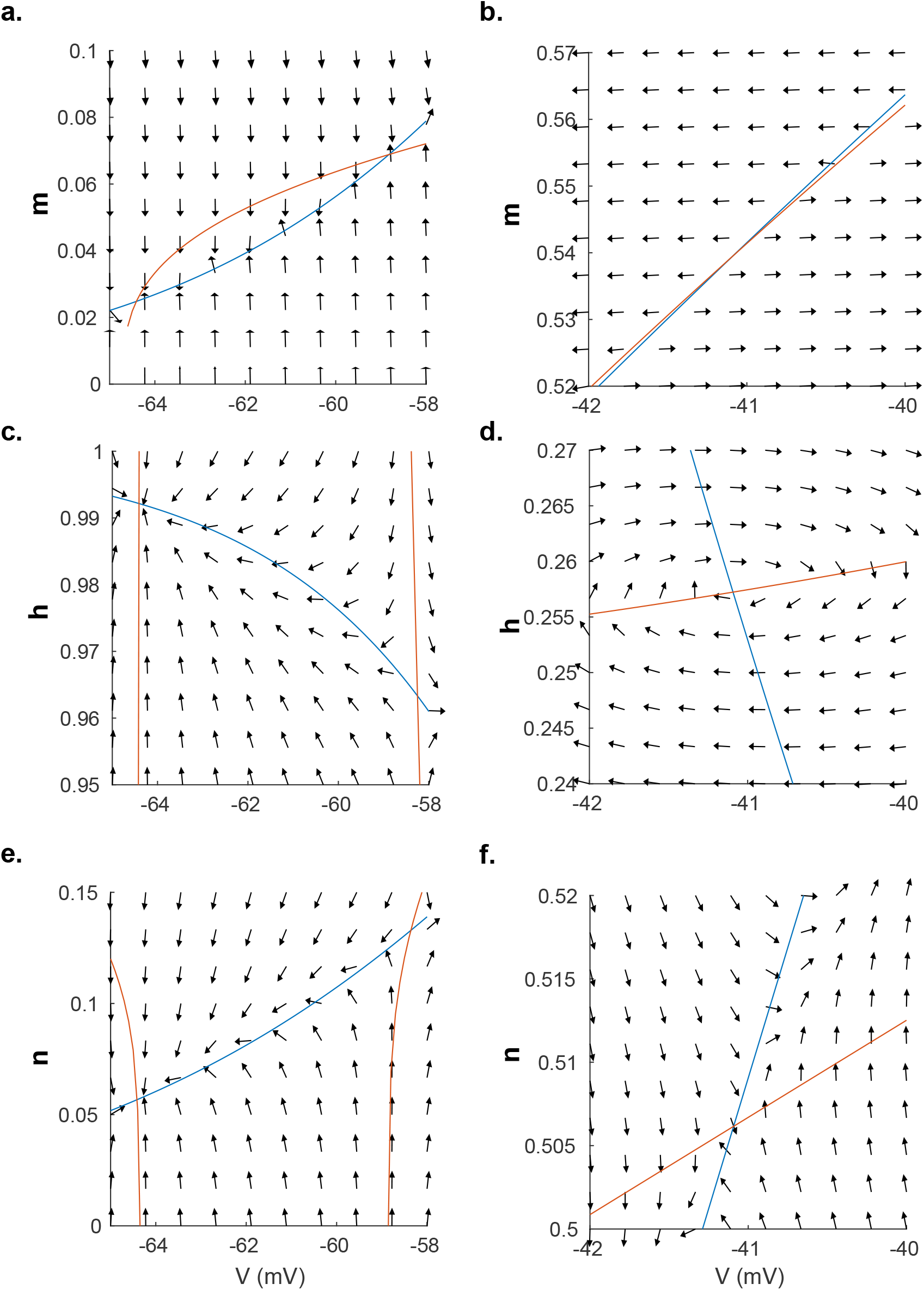
Field map around the equilibria that arises immediately following instantaneous pump rate increase by 10% for the three gating variables *m, n*, and *h* versus the membrane potential. Orange lines are the voltage nullclines (where 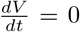), whereas the blue lines are gating variable nullclines (where 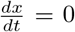 for *x* = *m, n, h*). The intersections of the nullclines are the equilibria. If the fields around the equilibrium all point towards it, it is a stable equilibrium, otherwise it is an unstable equilibrium. Only P1 near −64 mV has stable equilibrium for all three gating variables, showing that this is the only stable equilibrium in this system.

## Notes

### Competing Interest Statement

The authors have declared no competing interest.

